# Targeting glutamine metabolism enhances tumor specific immunity by inhibiting the generation of MDSCs and reprogramming tumor associated macrophages

**DOI:** 10.1101/656553

**Authors:** Min-Hee Oh, Im-Hong Sun, Liang Zhao, Robert Leone, Im-Meng Sun, Wei Xu, Samuel L. Collins, Ada J. Tam, Richard L. Blosser, Chirag H. Patel, Judson Englert, Matthew L. Arwood, Jiayu Wen, Yee Chan-Li, Pavel Majer, Rana Rais, Barbara S. Slusher, Maureen R. Horton, Jonathan D. Powell

## Abstract

Myeloid cells comprise a major component of the Tumor Microenvironment (TME) promoting tumor growth and immune evasion. By employing a novel small molecule inhibitor of glutamine metabolism not only were we able to inhibit tumor growth but we markedly inhibited the generation and recruitment of Myeloid Derived Suppressor Cells (MDSC). Targeting tumor glutamine metabolism led to a decrease in CSF-3 and hence recruitment of MDSC as well immunogenic cell death leading to an increase in inflammatory Tumor Associated Macrophages (TAMs). Alternatively, inhibiting glutamine metabolism of the MDSC themselves led to activation induced cell death and conversion of MDSC to inflammatory macrophages. Surprisingly, blocking glutamine metabolism also inhibited IDO expression of both the tumor and myeloid derived cells leading to a marked decrease in kynurenine levels. This in turn inhibited the development of metastasis and further enhanced anti-tumor immunity. Indeed, targeting glutamine metabolism rendered checkpoint blockade-resistant tumors susceptible to immunotherapy. Overall, our studies define an intimate interplay between the unique metabolism of tumors and the metabolism of suppressive immune cells.

## Introduction

The prodigious growth of tumor cells demands specialized metabolic reprogramming. Tumor metabolism not only promotes growth but also creates a TME that inhibits immune effector function by depleting critical metabolites (such as tryptophan, glucose and glutamine) and generating inhibitory metabolites such as kynurenine. Alternatively, suppressive immune cells, that are metabolically distinct from effector cells, thrive in the TME (Altman et al., 2016; DeBerardinis and Chandel, 2016; Pavlova and Thompson, 2016). To this end, the most prominent of immune cells in the TME are suppressive macrophages.

Macrophages, which constitute a major component of tumors, are involved in cancer initiation, progression, angiogenesis, metastasis, and creating an immune suppressive environment (Kondo et al., 2000; Mantovani et al., 2017; Noy and Pollard, 2014; Sica and Mantovani, 2012). Additionally, TAMs express enzymes like iNOS or arginase 1 (both enzymes that lead to arginine depletion) and IDO (an enzyme that leads to tryptophan depletion) that inhibits T cell activation and proliferation (Grivennikov et al., 2010; Kitowska et al., 2008; Lee et al., 2002; Mellor et al., 2002; Munn and Mellor, 2013). TAMs also express PDL1 and PDL2, which interact with PD1 on T cells (Rodriguez-Garcia et al., 2011). These interactions trigger inhibitory immune checkpoint signals on the T cells (Kryczek et al., 2006; Prima et al., 2017).

In addition to TAMs, MDSCs also play important roles in creating an immunosuppressive TME (Tcyganov et al., 2018). In mice, MDSCs express Gr1 (Ly6C and Ly6G) and CD11b. These markers define two subsets of MDSCs, polymorphonuclear-MDSCs (PMN-MDSCs, CD11b^+^ Ly6^lo^ Ly6G^+^) and monocytic-MDSCS (Mo-MDSC, CD11b^+^ Ly6C^hi^ Ly6G^-^). Though there are no distinct markers to distinguish between MDSCs and the tumor associated neutrophil (TAN)/monocytes at different stages of maturity, they are both functionally immunosuppressive cells in the TME (Bronte et al., 2000; Bronte et al., 2016; Li et al., 2004). Akin to TAMs, MDSCs also express enzymes that deplete key nutrients from T cells, express iNOS, arginase1, PDL1/2, and secrete suppressive cytokines (Schmielau and Finn, 2001; Serafini et al., 2008; Srivastava et al., 2010). Importantly, these cells do not highly express MHC and co-stimulatory molecules, which are essential for antigen presentation and activation to cytotoxic T cells (Almand et al., 2001).

In this report we employed a novel small molecule (Rais et al., 2016) to target glutamine metabolism. Our studies reveal that blocking glutamine metabolism markedly inhibits the generation and recruitment of MDSC and promotes the generation of anti-tumor inflammatory TAMs. Mechanistically we demonstrate a tumor specific and myeloid cell specific role for glutamine in promoting the immunosuppressive TME.

## Results

### Targeting glutamine metabolism inhibits tumor growth and MDSC recruitment in an immunotherapy-resistant model of triple negative breast cancer

The 4T1 triple negative breast cancer model is resistant to checkpoint blockade and this lack of response is associated with a low frequency of mutations and abundant presence of suppressive myeloid cells such as MDSCs, TAMs and TANs (Kim et al., 2014). This highly aggressive tumor model can form viable tumors when only 500 cells are implanted in mammary fat pad of female mice (Bailey-Downs et al., 2014; Gregorio et al., 2016; Pulaski and Ostrand-Rosenberg, 1998). In agreement with previous reports, 4T1 tumors were resistant to treatment with anti-PD1, anti-CTLA4, or combination of anti-PD1 and anti-CTLA4 (Figure 1A). 4T1 tumor-bearing mice showed elevated MDSCs in the blood compared to tumor free mice (Figure 1B). Treatment with immune checkpoint blockade had no impact on the recruitment of myeloid suppressor cells (Mo-MDSC : Live CD45^+^CD11b^+^ F4/80^neg^ Ly6C^hi^ Ly6G^neg^, and PMN-MDSCs and TANs (referred to as PMN-MDSCs due to a lack of markers to distinguish them : Live CD45^+^ CD11b^+^ F4/80^neg^ Ly6C^lo^ Ly6G^hi^) and CD8:MDSCs ratio in blood and in tumor (Figure 1B and C).

**Figure 1.**
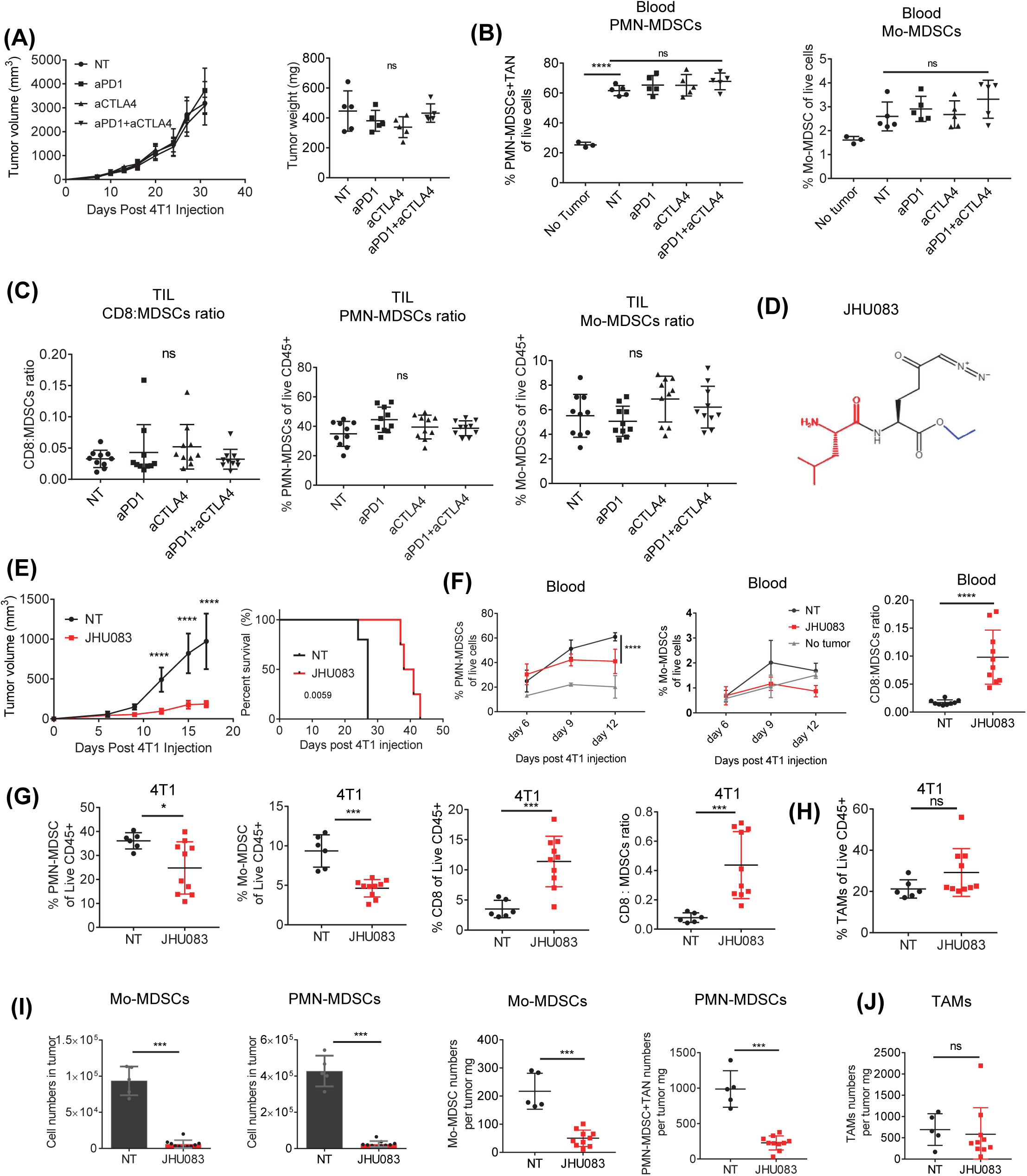
Glutamine antagonism inhibits tumor growth by enhancing CD8 population, and reducing MDSCs but not TAMs. 0.1×10^6^ 4T1 cells were implanted subcutaneously into the mammary fat pad of BALB/cJ female mice. On day 7, 10, 13, 17, and 24, mice were injected IP with 250 μg anti-PD1 and/or 100 μg anti-CTLA4 antibodies. (A) On day 17, percentages of PMN-MDSCs (CD11b^+^F4/80^neg^Ly6C^lo^Ly6G^hi^) and Mo-MDSCs (monocytic MDSCs: CD11b^+^F4/80^neg^Ly6C^hi^Ly6G^neg^) of live cells from the blood were analyzed by flow cytometry (N=5/group). (B) Tumor size was monitored (N=5/group) (left). On day 17, tumor weight was measured (right). (C) Ratio of CD8 cells to MDSCs, and percentages of PMN-MDSCs and Mo-MDSCs from the tumor were analyzed by flow cytometry (N=10/group). (D) The structure of the glutamine antagonist prodrug, JHU083. 6-Diazo-5-oxo-L-norleucine (DON: active glutamine antagonist) is depicted in black and its ethyl and 2-Amino-4-methylpentanamido promoieties are depicted in blue and red, respectively. 4T1 tumor-bearing mice were treated with JHU083 (1mg/kg) starting at day 7 after tumor inoculation. After 7 days of treatment, a lower dose (0.3 mg/kg) of JHU083 was used (E-J). (E) 4T1 Tumor sizes were measured and survival curve were recorded. (F) Percentages of PMN-MDSCs and Mo-MDSCs of live cells from blood in 4T1 tumor-bearing mice were analyzed by flow cytometry at the indicated time point (N=7-8/group). Ratio of CD8^+^ cells to MDSCs from blood was shown (N=9-10/group). (G) On day 14, tumors were harvested and tumor-infiltrating immune cells were analyzed by flow cytometry. The populations of PMN-MDSCs, Mo-MDSC, CD8^+^were shown. Ratio of CD8 cells to MDSCs in 4T1 tumor were evaluated. (H) Percentage of TAM (CD11b^+^F4/80^+^CD8^neg^Ly6c^neg^Ly6g^neg^) population among live CD45^+^ cells from 4T1 tumor-infiltrating immune cells (N=5-10/group). (I and J) On day 14, 4T1 tumors were harvested and each population cell numbers were counted. Total cell numbers were divided by respective tumor weights (mg). (N=5-10/group) Data are representative of at least three independent experiments or combined from two independent experiments (C). *P < 0.05, ***P < 0.005, ****P < 0.001, MEAN ± S.D. Kruskal-Wallis test with Dunn’s multiple comparisons post-test (A-C), Log-rank (Mantel-Cox) test (E) and Mann-Whitney t tests (F-J).

MDSCs themselves undergo metabolic reprogramming by increasing glycolysis, glutaminolysis, and fatty acid oxidation compared to mature granulocytes or monocytes (Hammami et al., 2012; Kumar et al., 2016; Li et al., 2018). This metabolic programming enables them to thrive in the harsh conditions of the TME (Gabrilovich, 2017; Sica and Strauss, 2017; Sieow et al., 2018). In light of the robust generation of MDSCs in the 4T1 model, we wanted to test the hypothesis that targeting glutamine metabolism might not only arrest tumor growth but also mitigate the generation and recruitment of these suppressive cells.

To this end, we employed a novel glutamine metabolism inhibitor prodrug of 6-Diazo-5-oxo-l-norleucine (DON) referred to as JHU083 (Figure 1D) (Rais et al., 2016). 4T1 tumor-bearing mice were treated with JHU083 (1 mg/kg) for 7 days starting at day 7 after tumor inoculation followed by a lower dose (0.3 mg/kg) until the mice were sacrificed. We observed a marked decrease in the growth of the 4T1 tumors following treatment with the glutamine antagonist, JHU083 (Figure 1E). We did not observe any weight loss due to our novel glutamine antagonist (Supplementary figure 1A). Interestingly, after 5 days of treatment we observed a marked reduction of PMN-MDSCs, and Mo-MDSCs in the blood compared to the control group (Figure 1F). More importantly, this decrease in MDSCs was associated with a slight increase in both the percentages of CD4^+^ and CD8^+^ T cells in the peripheral blood (Supplementary figure 1B). That is, the decrease in MDSC in peripheral blood was <u>not</u> due to a generalized decrease in total white blood cells. Along these lines, we observed a marked increase in the CD8:MDSCs ratio in blood from JHU083-treated mice (Figure 1F). Of note, a glutaminase specific inhibitor CB839 has been described (Gross et al., 2014; Wang et al., 2010; Xiang et al., 2015) and is currently undergoing clinical trials (Calithera Biosciences, 2014a, b, c). Using the established twice daily dosing regimens (from day 1 or 2 post tumor implantation), we did not observe any tumor growth delay in this 4T1 tumor model (data not shown). Thus, selective glutaminase inhibition is insufficient to inhibit tumor growth in the 4T1 model.

Next, we examined the effect of glutamine antagonism on tumor-infiltrating immune cells. Similar to what was observed in the blood, the percentages of both PMN-MDSCs and Mo-MDSCs were markedly reduced amongst the tumor-infiltrating immune cells of JHU083 treated tumor-bearing mice compared to the control group (Figure 1G). Additionally, we observed an increase in the tumor-infiltrating CD8^+^ T cells, and an enhanced ratio of CD8 to MDSCs ratio from JHU083 treated tumor-bearing mice compared to control group (Figure 1G). Interestingly, the percentage of TAMs (Live CD45^+^ CD11b^+^ F4/80^+^ Ly6C^neg^ CD8^neg^ Ly6G^neg^) was not different between the control and JHU083 treated group (Figure 1H). That is, treatment with JHU083 did not lead to a decrease in <u>all</u> myeloid cells within the TME, but rather led to the selective depletion of MDSCs. Concomitant with the reduction of the percentage of MDSCs within the tumors we also observed a decrease in the absolute numbers of these cells in the treated versus the untreated mice (Figure 1I). However, importantly the absolute number of TAMs per tumor weight did not change (Figure 1J). Finally, we tested the ability of glutamine antagonism to inhibit MDSCs in another immunotherapy resistant tumor model, the Lewis lung carcinoma (3LL) model (Supplementary Figure 1C). Similar to the 4T1 model, targeting glutamine metabolism in 3LL tumor-bearing mice led to improved control of tumor growth as well as a decrease in the percentage of MDSCs and an increase in the CD8:MDSCs ratio in blood and tumor (Supplementary Figure 1D). Overall these findings demonstrate that targeting glutamine metabolism not only inhibits tumor growth but also leads to a decrease in MDSCs.

### Targeting glutamine metabolism inhibits MDSCs by increasing cell death and decreasing tumor CSF3 expression

First, to understand glutamine antagonism effects on the decrease in MDSCs, we evaluated the direct effect of glutamine antagonism on MDSCs. When we treated MDSCs with DON (active form of JHU-083) in vitro, we observed an increase in apoptosis as defined by active caspase 3 (Figure 2A). Next, we evaluated the induction of MDSC apoptosis in vivo. To minimize the possibility that differences in tumor size itself affects MDSC numbers, we treated mice with JHU-083 for a short duration after the tumors were fully established (17 days after tumor inoculation). Consistent with our in-vitro finding, we also observed significantly increased active caspase 3 on both PMN-MDSCs and Mo-MDSCs within 33 hours after JHU-083 treatment in the blood (Figure 2B). Thus, these data suggest that glutamine antagonism directly affects apoptosis of MDSCs in the blood. Surprisingly, however, within 7hrs of treatment we observed markedly decreased tumor infiltrating MDSCs (Figure 2C). This finding led us to hypothesize in addition to inducing apoptosis in MDSC, blocking glutamine metabolism might indirectly affect the recruitment of MDSCs.

**Figure 2.**
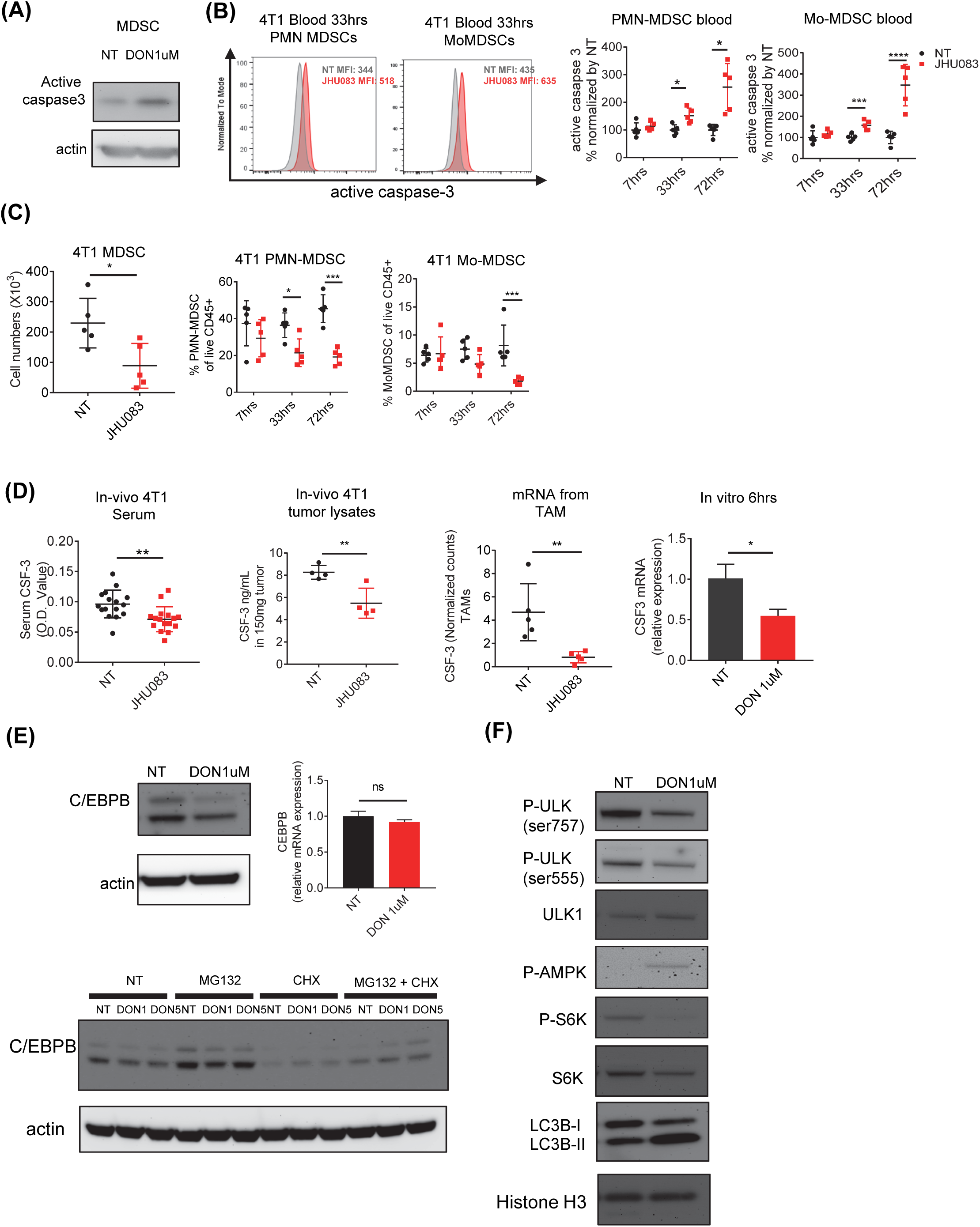
Glutamine antagonism reduces MDSCs by increased cell death and inhibition of tumor CSF3 secretion. (A) MDSCs from 4T1 tumor-bearing mice were treated with DON1µM for 24hrs, and active caspase-3 levels was analyzed by immunoblot. β-actin was used as loading control. 4T1 tumor-bearing mice were treated with JHU083 (1 mg/kg) starting at day 14 after tumor inoculation. After the indicated hours of treatment, (B) active caspase 3 on PMN-MDSCs and Mo-MDSCs from blood (C) Cell numbers of MDSCs, percentages of PMN-MDSCs and Mo-MDSCs, and active caspase 3 from tumors were analyzed by flow cytometry. (D) Serum and tumor were collected from 4T1 tumor-bearing mice treated with or without JHU083 on day 21. CSF-3 was measured by ELISA (N=16/group). Normalized transcript counts of CSF-3 by RNA sequencing on TAMs from NT and JHU083-treated mice. (N=5/group, q-value <0.05). 4T1 tumor cells were treated with DON1µM (D-F). After 6hrs treatment, (D) CSF3 and (E) After 24hrs alter, CEBPB protein levels were measured by immunoblotting (left). CEBPB mRNA levels were measured by q-PCR (right). 4T1 cells were treated with or without DON1 or 5µM in the presence or absence of MG-132 (1µM) or cyclohexamide (20 µM) for 4 hrs. CEBPB were analyzed by immunoblotting. (F) After treatment with DON1µM for 24hrs, autophagy related proteins were measured by immunoblotting. MEAN ± S.D. Two-way ANOVA with post multiple t tests (B and C) and Mann-Whitney t tests (D and E).

In this regard, several studies have demonstrated that increased secretion of growth factors such as CSF-1, CSF-2, and CSF-3 promote the recruitment of MDSCs to the TME (Condamine et al., 2015). As such, we wanted to determine if targeting glutamine metabolism inhibited MDSCs in the TME in part by limiting the elaboration of these critical growth factors. To this end we measured CSF-3 (G-CSF) levels in the serum of 4T1 tumor-bearing mice treated with or without JHU083. Compared to the vehicle group, the glutamine antagonist treated mice demonstrated reduced CSF-3 in circulating serum (Figure 2D). Furthermore, we detected markedly decreased CSF-3 protein in tumor lysates, and CSF3 mRNA expression from the TAMs isolated from the JHU083 treated mice. Additionally, in-vitro DON treated 4T1 cells also showed reduced CSF3 mRNA expression (Figure 2D). Such findings support the notion that one mechanism by which glutamine antagonism leads to a decrease in MDSCs within the TME is through inhibiting the transcriptional expression of CSF-3 from tumor cells and TAMs. To dissect the mechanism of transcriptional regulation of CSF-3 in tumor cells, we evaluated the expression of C/EBPB, a well described transcription factor for CSF-3 expression (Akagi et al., 2008; Li et al., 2018; Marigo et al., 2010). Interestingly, even though we observed markedly decreased C/EBPB protein expression, we did not observe any mRNA expression differences of C/EBPB in the presence or absence of DON treatment (Figure 2E). Next, we wondered whether glutamine antagonism controls the stability of C/EBPB by regulating protein degradation. To address this question, we treated cells with DON in the presence or absence of the proteosomal degradation inhibitor, MG132, and/or the translation inhibitor, cyclohexamide. Indeed, translation of C/EBPB protein was markedly reduced by treatment with cyclohexamide. On the contrary, treatment with MG132 or MG132+cyclohexamide enhanced the expression of C/EBPB protein on DON treated 4T1 cells suggesting that protein degradation actively regulates protein levels.

A previous report has shown that C/EBPB is the target for protein degradation through the autophagy process (Li et al., 2018). Since glutamine metabolism is closely related to AMPK and mTOR signaling which are important in regulating autophagy, we thought altered AMPK and mTOR activity via glutamine metabolism inhibition might enhance autophagy, and sequentially, induce C/EBPB degradation. As expected, increased AMPK and reduced mTOR activity were observed in DON treated 4T1 tumor cells (Figure 2F). Accordingly, phosphorylation of ULK1, which is directly regulated by AMPK and mTOR, and are important in initiation of autophagosome formation, was reduced in DON treated cells. As a result, autophagy flux measured by LC3BI to LC3II was increased in DON treated cells. Thus, these data suggest that glutamine metabolism is important for the maintenance of C/EBPB, which is crucial for MDSCs recruitment via CSF3 expression. Overall our data thus far demonstrate that targeting glutamine metabolism, in addition to inhibiting tumor growth, has a robust effect on inhibiting MDSCs in immunotherapy resistant tumors. Mechanistically this is through two distinct effects: i: Direct effects on MDSC promoting caspase-3 dependent cell death. ii. Effects on the tumor by inhibiting the elaboration CSF-3 mechanistically by promoting autophagy thereby inhibiting C/EBPB transcription factor activity.

### Targeting glutamine metabolism promotes the reprogramming of tumor associated macrophages

While targeting glutamine metabolism inhibited the recruitment of Mo-MDSCs and PMN-MDSCs to the tumor, it did not “wholesale” inhibit myeloid cells. Recall, we did not observe significant differences in the percentage of TAMs in the tumors from the treated and untreated mice (Figure 1H and J). Thus, we were interested in understanding the effect of JHU083 on the phenotype and function of the TAMs. To this end, we performed RNA sequencing on sorted TAMs from vehicle and JHU083-treated 4T1 tumor-bearing mice. Distinct transcriptional changes between these two groups were observed and more than 3,000 significant mRNA transcripts were differentially expressed (Figure 3A). As expected, we found significant differences between TAMs from vehicle and JHU083 treated mice within glutamine related pathways, such as DNA replication, cell cycle, pentose phosphate pathway, glycolysis, pyrimidine and purine metabolism, and arginine and proline metabolism (Table 1).

**Figure 3.**
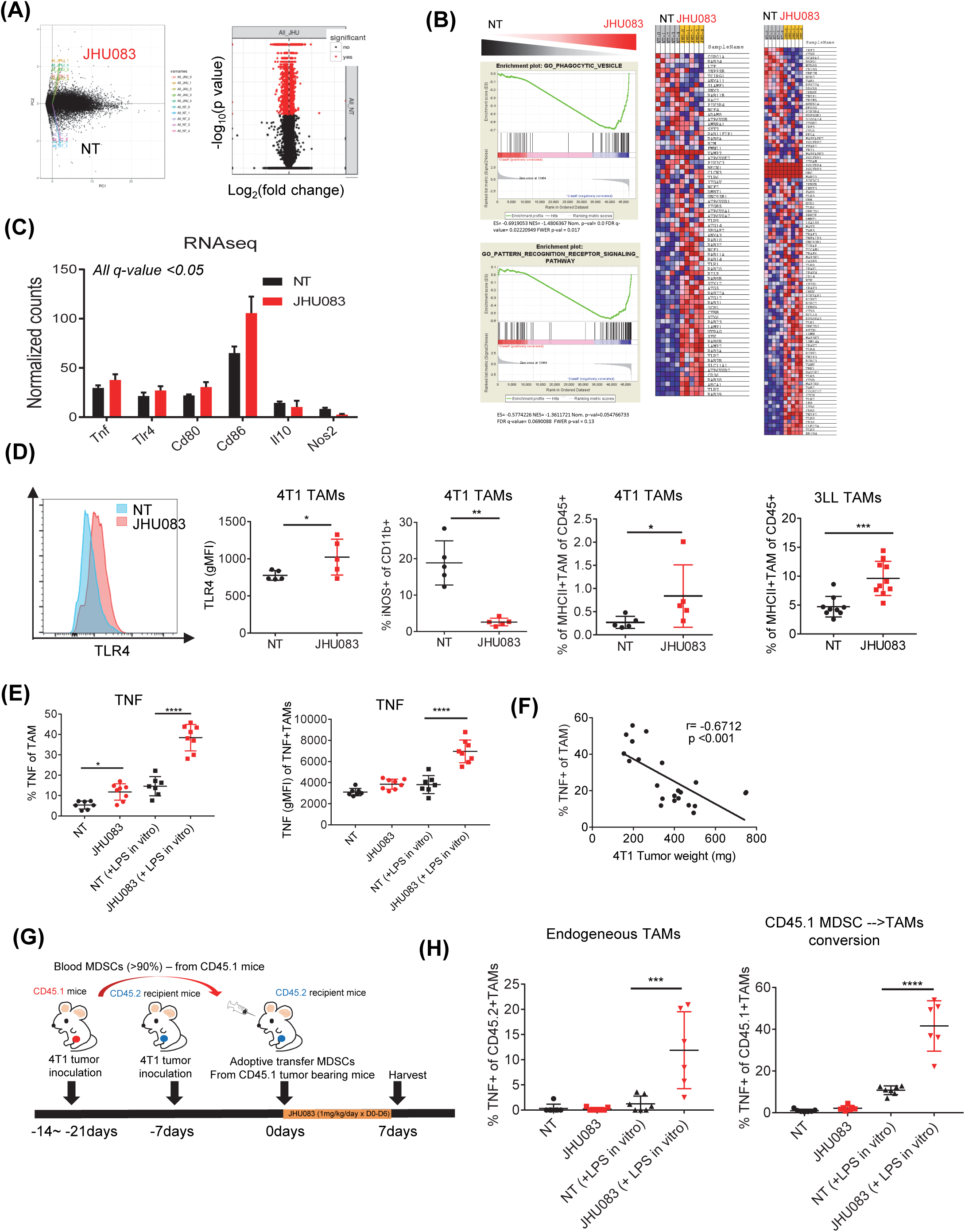
Glutamine antagonism induces reprogramming of TAMs and differentiation of MDSCs from suppressive to a pro-inflammatory phenotype. (A) Principal component analysis and volcano plot showing changes in gene expression (red) from RNA sequencing analysis on NT and JHU083 treated TAMs (CD11b^+^F4/80^+^7AAD^neg^Ly6C^neg^Ly6G^neg^CD8^neg^) from 4T1 tumor-bearing mice (on day 14). q-value <0.05 (B) Gene set enrichment analysis (GSEA) plot of phagocytic vesicle and signaling pattern recognition receptor activity related genes in NT vs. JHU083 on TAMs. Enrichment scores (ES), gene set null distribution, and heat map for genes in gene set are shown. (C) Normalized gene expression from RNA sequencing analysis on NT (black) and JHU083 (red) treated TAMs from 4T1 tumor-bearing mice (on day 17). All genes are significant (q-value <0.05). (D) Representative histogram of TLR4 expression on TAMs. Summary graph of TLR4 and iNOS expression. Percentage of MHCII^+^ TAMs from 4T1 tumor-bearing mice and 3LL tumor-bearing mice were analyzed by flow cytometry. (E) TILs were harvested on day 17 from 4T1 tumor-bearing mice treated with or without JHU083. Cells were incubated with golgi-plug in the presence or absence of LPS for 9 hours ex-vivo. Percentages of TNF^+^ cells were analyzed by flow cytometry (left). MFI of TNF from TNF^+^ cells (right). (F) Correlation of % of TNF^+^ secreting TAMs after stimulation with respective to tumor weight. Isolated MDSCs in blood from CD45.1 4T1 tumor bearing mice (21 days after 4T1 tumor inoculation) were adoptively transferred into CD45.2 4T1 tumor bearing mice (7days after 4T1 tumor inoculation). Then, MDSCs transferred CD45.2 4T1 tumor bearing mice were treated with JHU083 until harvesting tumors on day 7. (G) Schematic of the experiment (H) Cells were incubated with golgi-plug in the presence or absence of LPS for 9 hours ex-vivo. Percentages of TNF^+^ CD45.1^+^ cells (adoptive transferred, left) and CD45.2^+^ cells (endogenous, right) were analyzed by flow cytometry. Data are from one experiment with 5 mice per group (A-C) or from three independent experiment with 5-10 mice per group (D-H) *P < 0.05, **P < 0.01, ***P<0.005, ****P<0.001, MEAN ± S.D. Mann-Whitney t tests (D and E) and spearman correlation (F).

**Table 1.** Gene ontology analysis of RNA sequencing data on sorted TAMs from WT and JHU083 treated 4T1 tumor-bearing mice. Molecular functional analysis using gene ontology in up-regulated genes and down regulated genes (q value < 0.05)

**GO molecular function analysis in up-regulated genes in JHU083 treated TAMs**
| Term | Count | % | P-Value |
| --- | --- | --- | --- |
| Lysosome | 39 | 0.2 | 2.9E-14 |
| Toll-like receptor signaling pathway | 22 | 0.1 | 0.000031 |
| Sphingolipid metabolism | 12 | 0.1 | 0.00037 |
| Endocytosis | 32 | 0.2 | 0.00034 |
| Hematopoietic cell lineage | 18 | 0.1 | 0.00032 |
| Chemokine signaling pathway | 28 | 0.2 | 0.0014 |
| Circadian rhythm | 6 | 0 | 0.0025 |
| Complement and coagulation cascades | 15 | 0.1 | 0.0024 |
| MAPK signaling pathway | 36 | 0.2 | 0.0023 |
| Renal cell carcinoma | 14 | 0.1 | 0.0035 |
| Cytokine-cytokine receptor interaction | 33 | 0.2 | 0.004 |
| Focal adhesion | 27 | 0.2 | 0.0088 |
| GnRH signaling pathway | 16 | 0.1 | 0.011 |
| TGF-beta signaling pathway | 14 | 0.1 | 0.022 |
| Fructose and mannose metabolism | 8 | 0 | 0.027 |
| Type II diabetes mellitus | 9 | 0.1 | 0.041 |

**GO molecular function analysis in down-regulated genes in JHU083 treated TAMs**
| Term | Count | % | P-Value |
| --- | --- | --- | --- |
| DNA replication | 22 | 0.1 | 6.7E-13 |
| Cell cycle | 42 | 0.2 | 7.2E-12 |
| Proteasome | 19 | 0.1 | 0.00000031 |
| Spliceosome | 32 | 0.2 | 0.0000015 |
| Pyrimidine metabolism | 26 | 0.1 | 0.0000072 |
| ECM-receptor interaction | 23 | 0.1 | 0.000019 |
| Pentose phosphate pathway | 11 | 0.1 | 0.00014 |
| Aminoacyl-tRNA biosynthesis | 14 | 0.1 | 0.00018 |
| One carbon pool by folate | 8 | 0 | 0.00057 |
| Mismatch repair | 8 | 0 | 0.0049 |
| Base excision repair | 11 | 0.1 | 0.0058 |
| Glycolysis / Gluconeogenesis | 15 | 0.1 | 0.0078 |
| Purine metabolism | 27 | 0.2 | 0.0087 |
| Nucleotide excision repair | 11 | 0.1 | 0.0099 |
| Terpenoid backbone biosynthesis | 6 | 0 | 0.01 |
| Small cell lung cancer | 17 | 0.1 | 0.011 |
| Oocyte meiosis | 21 | 0.1 | 0.012 |
| Galactose metabolism | 8 | 0 | 0.016 |
| RNA degradation | 13 | 0.1 | 0.017 |
| Glyoxylate and dicarboxylate metabolism | 6 | 0 | 0.018 |
| Steroid biosynthesis | 6 | 0 | 0.024 |
| Parkinson's disease | 22 | 0.1 | 0.028 |
| Arginine and proline metabolism | 11 | 0.1 | 0.04 |

Notably, by evaluating pathway analysis for biological processes, we found that lysosome and TLR signaling related genes significantly differed between TAMs isolated from tumors from JHU083 treated and untreated mice (Table 1). Furthermore, using gene set enrichment analysis (GSEA), we found an upregulation of phagocytic vesicles and signaling pattern recognition receptor activity related genes in TAMs from the treated mice (Figure 3B). Specifically, the expression of genes encoding for TNF, TLR4, CD80, and CD86 - molecules related to a pro-inflammatory phenotype were increased while *Nos2* and *Il10* gene expression, which is known to inhibit anti-tumor T cell responses, were decreased (Figure 3C). To confirm our RNA sequencing data, we performed flow cytometry to analyze the TAM phenotypes within the TME in 4T1 tumor-bearing mice. We observed increased surface TLR4 and MHCII, and reduced iNOS on TAMs from the JHU083 treated mice (Figure 3D). Similarly, we also found increased MHCII expression on TAMs from JHU083 treated 3LL-tumor bearing mice (Figure 3D). Overall, our findings demonstrate that targeting glutamine metabolism promotes a pro-inflammatory phenotype amongst TAMs.

Recently, multiple studies have demonstrated that pro-inflammatory TAMs inhibit tumor growth (Hoves et al., 2018; Perry et al., 2018). Our RNA sequencing data of TAMs from JHU083 treated mice demonstrated an increase in the pro-inflammatory cytokine, *Tnf*. In agreement with the RNA seq data, we also observed increased TNF protein production in TAMs from the treated mice (Figure 3E). After in vitro LPS stimulation for 9 hours, further enhancement of TNF production was also observed in TAMs from JHU083 treated mice compared to TAMs from vehicle treated mice (Figure 3E). Furthermore, there was a negative linear relationship between TNF production and tumor weight (Figure 3F). Consistent with our findings, a previous report demonstrated that glutamine deprivation further induces M1 polarization mediated by increased NF-κB signaling (Liu et al., 2017).

To confirm this finding, we treated bone marrow-derived macrophages (BMDMs) with varying doses of DON during LPS stimulation. After 24 hours, we observed increased TNF secretion with DON treatment in BMDMs along with increased NF-κB nuclear localization (Supplementary Figure 2A and B). On the other hand, we observed decreased IL-10 secretion and phosphorylation of STAT3. (Supplementary Figure 2C and D). Similarly, we observed a dose dependent increase in TNF production from DON treated BMDMs using flow cytometry analysis (Supplementary Figure 2E). Thus, this finding confirms that glutamine inhibition enhances a pro-inflammatory macrophage phenotype. Mechanistically this is due to increased NF-κB and reduced STAT3 signaling.

Given the observation that glutamine antagonism promotes the reprogramming of tumor associated macrophages, we hypothesized that recruited MDSCs in the tumor might be converted into pro-inflammatory macrophages. To this end, isolated MDSCs in the blood from CD45.1 4T1 tumor bearing mice (21 days after 4T1 tumor inoculation) were adoptively transferred into CD45.2 4T1 tumor bearing mice (7days after 4T1 tumor inoculation). Then, MDSCs transferred CD45.2 4T1 tumor bearing mice were treated with JHU083 until harvesting tumors on day 7 (Figure 3G). As seen in figure 4E, we observed increased TNF secreting endogenous TAMs in JHU083 treated mice. More strikingly, we found significantly increased TNF production from the adoptively transferred CD45.1^+^cells in tumor from JHU083 treated CD45.2 mice (Figure 3H). That is, adoptively transferred MDSC were converted to inflammatory TAMs upon treatment with JHU-083. These observations support a model whereby glutamine antagonism enhances pro-inflammatory TAM differentiation in not only TAMs but also in MDSCs.

**Figure 4.**
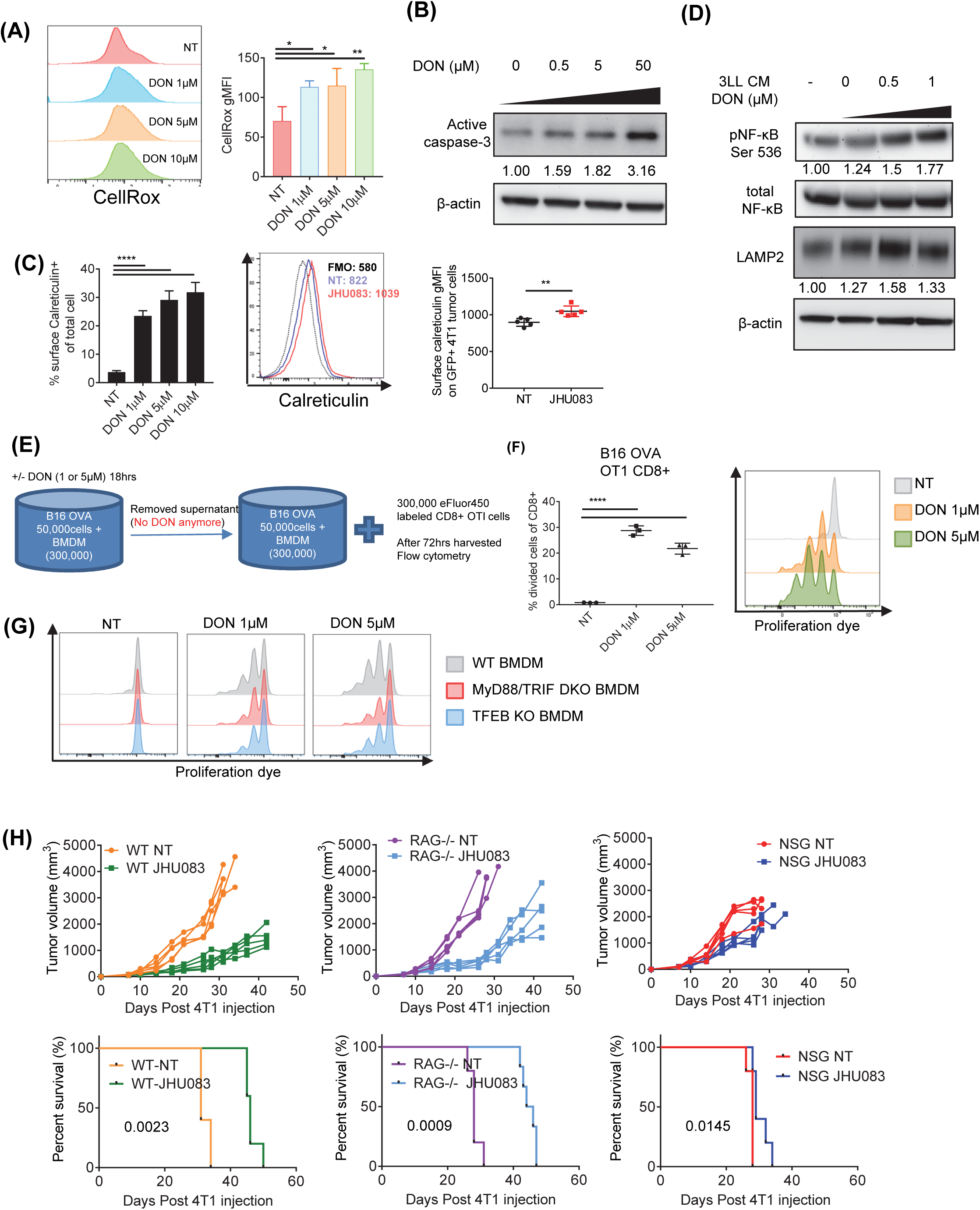
Glutamine antagonism increases immunogenic cell death and antigen presentation of macrophages to T cells. (A-C) 4T1 tumor cells were cultured with or without DON (0, 0.5 μM, 1 μM, 5 μM, 10 μM) for 24 hours. (A) Cells were harvested and stained with CellROX (ROS measurement), and analyzed by flow cytometry. Representative histogram (left) and summary graph (right). (B) Cells were lysed and active caspase-3 levels was analyzed by immunoblot. β-actin was used as loading control. (C) Cells were stained for calreticulin and analyzed by flow cytometry. Percentages of surface calreticulin were shown (Left). Representative histogram (middle) and summary graph of surface calreticulin gMFI on GFP^+^ gated tumor cells (Right). (D) 3LL cells were cultured in the presence or absence of DON (0.5 μM or 1 μM). After 1 hour of incubation, cells were washed and replaced with drug-free fresh media. After 24 hours, supernatants were harvested and used as conditioned media (CM). BMDMs were cultured in the presence of these conditioned media for 24 hours. Phospho-NF-κB (ser536) and LAMP2 were measured by immunoblot. Total NF-κB and β-actin were measured as loading controls. (E-G) 0.3×10^6^ BMDMs and 5×10^4^ B16-OVA tumor cells were co-cultured in the presence or absence of 1 μM or 5 μM of DON. After 24 hours of incubation, supernatants were discarded and 0.3×10 eFluor450-labeled CD8^+^ from OTI mice were added. Percentages of divided cells from CD8^+^ population were analyzed by flow cytometry. (E) Schematic of the experiment. (F) Percentages of divided cells from CD8^+^ population were analyzed by flow cytometry (left). Histogram showing the dilution of eFluor450-labeled CD8^+^ cells (right). (G) 0.3×10^6^ BMDMs from WT, MyD88/TRIF double KO or TFEB KO mice and 5×10^4^ B16-OVA tumor cells were co-cultured in the same method as (E). (F-G) Histogram showing the dilution of eFluor450-labeled CD8^+^ cells. (H) 0.1×10^6^ 4T1 cells were implanted subcutaneously into the mammary fat pad in NSG or RAG1 or BALB/cJ female mice. 4T1 tumor-bearing mice were treated with JHU083 (1mg/kg) daily starting at day 7 after tumor inoculation. After 7 days of treatment, a lower dose (0.3 mg/kg) of JHU083 was used. Tumor burdens were assessed. Data are representative of two (E) or three independent experiments (A-D). **P < 0.01, ***P<0.005, MEAN ± S.D. Mann-Whitney t tests (C). Two-way ANOVA with post multiple t tests (A-F). Log-rank (Mantel-Cox) test (H).

### Glutamine antagonism enhances immunogenic tumor cell death

Though we observed intrinsic enhancement of pro-inflammatory macrophage phenotypes with glutamine inhibition upon LPS stimulation, it is unclear how TAMs were activated in the TME with glutamine antagonist treatment without LPS stimulation. Previous reports have shown that immunogenic tumor cell death (ICD) induces TLR signaling in TAMs through the release of Danger-Associated Molecular Patterns (DAMPs) (Green et al., 2009). Increased endoplasmic reticulum stress and reactive oxygen species (ROS) production are important mediators in inducing ICD (Krysko et al., 2012). Thus, we investigated the ability of JHU083 to promote a pro-inflammatory TME by inducing ICD. Indeed, treatment of 4T1 cells with DON led to an increase in ROS and active-caspase 3 (Figure 4A and B). In addition, targeting glutamine metabolism of 4T1 tumor cells both in vitro and in vivo led to an increase in surface exposure of calreticulin, a DAMP and endoplasmic reticulum protein (Figure 4C). To explore this concept further, we cultured BMDM cells in conditioned media from DON-treated tumor cell supernatants. Increased p-NF-κB (ser536) (TLR downstream signaling) and LAMP2 (lysosome function marker) were observed in BMDMs cultured in DON-treated tumor conditioned media compared to vehicle treated tumor conditioned media (Figure 4D). This result suggests that tumor cell death induces macrophage activation mediated by downstream TLR signaling and lysosome function, which correlated with the RNA sequencing data (Table1).

Next, we tested whether the increased NF-κB signaling and lysosome function by ICD indeed increased antigen presentation to T cells. To test this idea, BMDMs were co-cultured with the B16 OVA melanoma tumor with various doses of DON treatment for 24 hours. After removing and washing away the media, cell proliferation dye-labeled naïve CD8^+^ T cells from OTI mice were co-cultured with BMDMs and tumor cells (Figure 4E). Next, dividing OVA-specific cytotoxic T cell populations were analyzed by flow cytometry. When compared to the vehicle-treated group, BMDMs co-cultured with DON-treated tumors showed increased OVA-specific T cell proliferation (Figure 4F). This finding also held true using the MC38 OVA tumor (Supplementary Figure 3A and B). When we added the drug to the macrophages prior to the co-culture, we did not observe proliferating CD8 T cells despite increased MHCII expression with glutamine inhibitor treatment (Supplementary Figure 3C). This finding demonstrates that macrophages require danger signals to trigger proper antigen presentation to T cells despite the enhanced activation itself with DON treatment. On the other hand, when we added DON to the tumor cells prior to the co-culture, we observed increased proliferating CD8 T cells. However, it was less effective compared to the co-culture of macrophages and tumor cells with drug treatment. Thus, glutamine inhibition exerts a synergistic effect between immunogenic tumor cell death and enhanced activation of macrophages by glutamine inhibition itself (Supplementary Figure 3C). Furthermore, when we co-cultured T cells with BMDMs from MyD88/TRIF (downstream of TLR signaling) or TFEB (key regulator of lysosomal biogenesis) deficient mice, there was diminished T cell proliferation (Figure 4G).

Thus, DON-induced tumor cell death enhances antigen presentation (as determined by T cell proliferation) in a MyD88/TRIF signaling and lysosome dependent fashion. Taken together, these data demonstrate that glutamine inhibition induces ICD of tumor cells by increasing ROS which leads to an increase in MyD88/TRIF-dependent signaling, lysosomal function, and antigen presentation in TAMs, and subsequently enhanced tumor-specific effector T cell proliferation.

To dissect the contribution of immune cells in the reduction of tumor size in JHU083 treated mice 4T1 tumors were injected into mammary fat pad of non-obese diabetic severely combined immune-deficient interleukin-2 receptor gamma-chain null (NSG) mice lacking adaptive immunity with <u>defective innate immunity,</u> or RAG1 KO mice lacking adaptive immunity with <u>intact innate immunity</u> or immunocompetent wild type mice (WT). JHU083 treated NSG mice showed minimal therapeutic effects compared to JHU083 treated RAG1 KO and WT mice (Figure 4H). Given the fact that NSG mice have defective macrophages and that macrophages are a major component of the TME, these data are consistent with the profound effect of JHU083 treatment on TAM reprogramming (Figure 3). Furthermore, JHU083 treated RAG1 KO mice controlled tumor size just as well as JHU083 treated WT mice in the early phase of tumor growth, confirming the effects of reduced suppressive myeloid cells and activation of TAMs. However, in the late phase of tumor growth, JHU083 treated RAG1 KO mice showed faster tumor growth compared to JHU083 treated WT indicating the importance of adaptive immune responses against tumor with the therapeutic effects of JHU083 (Figure 4H). Altogether, these data suggest that JHU083 triggers immune responses against tumor through enhanced innate immunity that sequentially elicits adaptive immune responses.

### Targeting glutamine metabolism alters the metabolism of the tumor in both glutamine dependent and glutamine independent pathways

Glutamine is a critical anaplerotic substrate for anabolic growth that is necessary for the specialized Warburg metabolism that facilitates robust tumor growth (Altman et al., 2016; Yang et al., 2017). We hypothesized that blocking glutamine metabolism would not only inhibit tumor growth but also alter the metabolic milieu of the TME. To this end, we performed targeted metabolomics using liquid-chromatography tandem mass spectrometry (LC-MS/MS) on equal weighted 4T1 tumors from vehicle and JHU083-treated mice to assess the effects of glutamine inhibition on cell metabolism. Metabolomics analysis with LC-MS revealed two distinct metabolic clusters, which correlated to the two experimental groups (Figure 5A and 5B). As expected, glutamine inhibition caused reduced TCA cycle intermediates and less conversion from glutamine to glutamate, resulting in an increased glutamine/glutamate ratio, implying GLS inhibition (Supplementary Figure 4A). In addition, we also observed significant changes in citrulline, N-carbamoyl-L-aspartate, thymine, S-adenosyl-L-methionine, homoserine, guanosine, nicotinamide ribotide, hydroxyproline, succinate, cystathionine, aspartate, uridine, Acetyl-L-lysine, and Dimethyl-L-arginine. Using pathway enrichment analysis, we found significant differences between the metabolites in tumors from vehicle and JHU083 treated mice, such as in glycine and serine metabolism, phosphatidylcholine biosynthesis, methionine metabolism, urea cycle, glutamate metabolism, ammonia recycling, amino sugar metabolism, and arginine and proline metabolism pathways. (Figure 5B and Supplementary Figure 4A).

**Figure 5.**
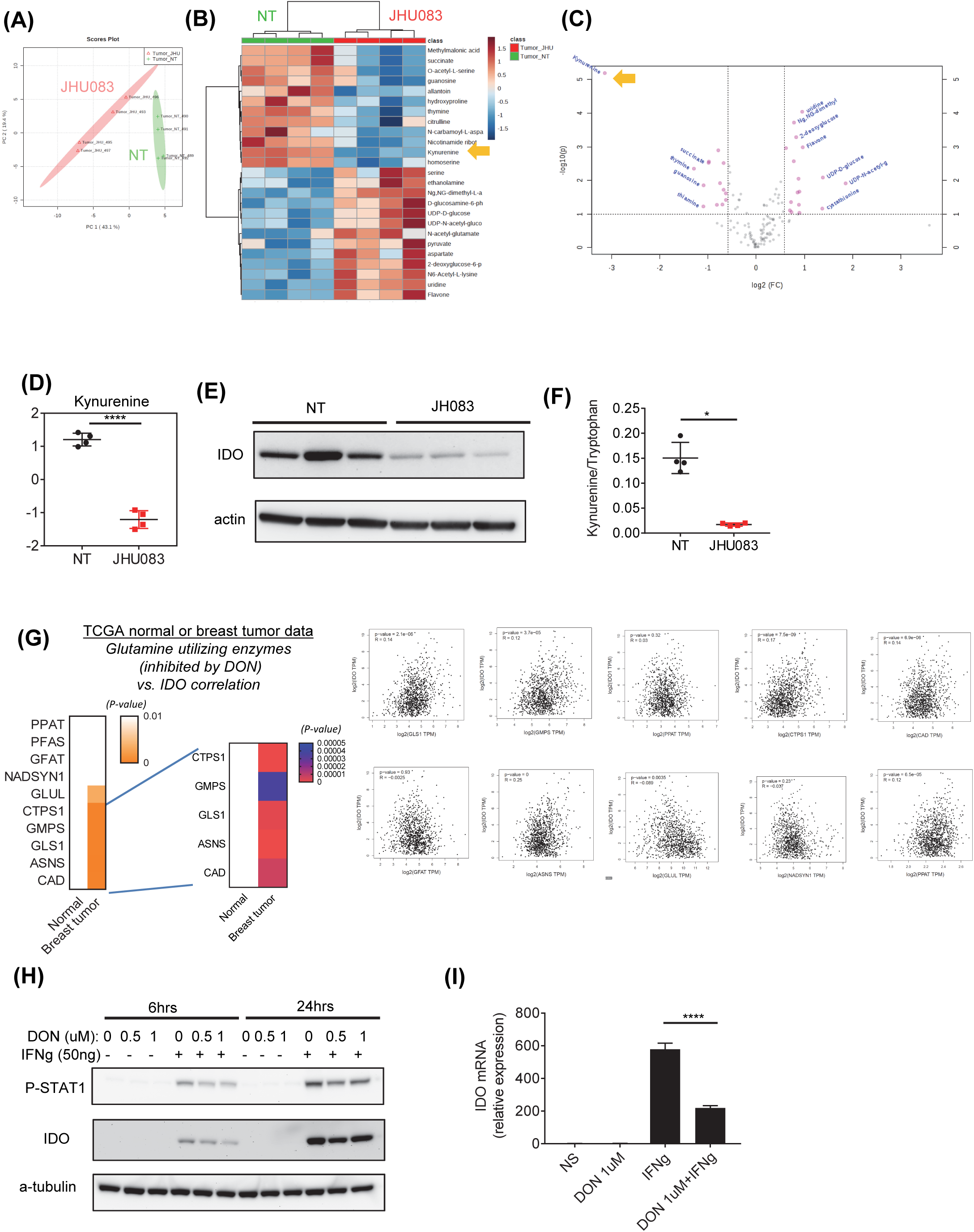
Glutamine antagonism alters primary tumor metabolism in both glutamine dependent and independent pathways. 0.1×10^6^ 4T1 cells were implanted subcutaneously into mammary fat pad of BALB/cJ female mice. 4T1 tumor-bearing mice were treated with JHU083 (1 mg/kg) starting at day 7 after tumor inoculation. After 7 days of treatment, a lower dose (0.3 mg/kg) of JHU083 was used. On day 17, tumors were harvested, and 95 mg of tumor samples were applied for LC-MS analysis. (A) Principal component analysis between NT (vehicle treated, green) and JHU083-treated (red) groups, (B) heatmap visualization of the metabolite changes between NT (green) and JHU083-treated (red) groups, and (C) volcano plot of metabolites were shown. Log2 fold change vs. –log10 (FDR corrected P value) representing significant metabolite changes. Red: significant. (D) Relative amounts of kynurenine between NT and JHU083-treated group. (E) On day 21, IDO expression in tumor lysates from 4T1 tumor-bearing mice were measured by immunoblot. β -actin was used as loading control. (F) The ratio of kynurenine to tryptophan in tumor was shown. (G) Heat map visualization of p-value from pearson correlation analysis (non-log scale for calculation) using TCGA normal and breast invasive carcinoma data between the glutamine utilizing enzymes which are inhibited by DON and IDO expression. The p-value close to 0 is more significant compared to 1 (left). The graphs from each enzyme and IDO correlation data. (H) 4T1 cells were cultured in the presence or absence of DON (0.5 μM or 1 μM) for 6 or 24 hours. P-STAT1 (ser S727) and IDO were measured by immunoblot. α-tubulin were measured as loading control. (I) After 6hrs with DON 1 μM treatment, IDO mRNA expression level was measured by q-PCR. *P < 0.05, ****P<0.001, MEAN ± S.D. t-test (D) and Mann-Whitney t tests (F). Two-way ANOVA (I).

Interestingly, of the 200 molecules tested, kynurenine was found to be the most reduced metabolite in the treated mice (Figure 5C and D). This was quite surprising since glutamine is not known to be directly involved in kynurenine metabolism. Rather, the enzyme IDO metabolizes tryptophan to kynurenine which is a potent inhibitor of T cell proliferation and function (Munn and Mellor, 2016). Thus, we hypothesized that glutamine antagonism regulates IDO expression but not its enzymatic activity. Indeed, IDO protein expression in the tumor lysates was decreased in the JHU083 treated mice compared to vehicle treated mice (Figure 5E). To determine the major source of IDO in tumor, we performed immunoblotting on different cell populations in the TME. We observed that tumor cells are the major source of IDO and JHU083 treatment dramatically decreases IDO expression (Supplementary Figure 4B). Additionally, sorted MDSCs and TAMs express IDO that is also markedly reduced with JHU083 treatment (Supplementary Figure 4B). Furthermore, the ratio of kynurenine: tryptophan was markedly diminished in the tumors from the JHU083 treated mice (Figure 5F). Interestingly, analysis of The Cancer Genome Atlas using normal and breast invasive carcinoma data revealed significant correlation between expression levels of IDO and glutamine utilizing enzymes which are inhibited by DON in breast carcinoma samples but not normal tissue controls (Figure 5G).

Furthermore, IDO is known to be transcriptionally regulated by STAT1 and STAT3. To understand how IDO expression is regulated by glutamine metabolism, we measured the phosphorylation of STAT1 as an indicator of its transcriptional activity. With DON treatment, we observed reduced p-STAT1 in tumor along with reduced IDO expression upon IFN gamma stimulation (Figure 5H). Accordingly, reduced mRNA expression of IDO was observed with glutamine inhibition (Figure 5I). Similarly, with DON treatment, we observed reduced p-STAT3 in macrophages along with decreased IDO protein and mRNA expression upon IFN gamma stimulation (Supplementary figure 4C). Thus, targeting glutamine metabolism not only inhibits glutamine dependent pathways, but also somewhat surprisingly, leads to a marked decrease in p-STAT1 and p-STAT3 dependent IDO expression resulting in a robust reversal of kynurenine: tryptophan ratio.

### Targeting glutamine metabolism inhibits lung metastasis by altering metabolism at the site of metastasis

The 4T1 tumor model is known to have high metastatic potential. Interestingly, both MDSC and the metabolite kynurenine have been implicated in promoting metastasis ((D’Amato et al., 2015; Safarzadeh et al., 2018; Smith et al., 2012; Wang et al., 2019; Xue et al., 2018). Thus, we wondered whether glutamine inhibition could prevent metastasis. Indeed, in addition to inhibiting the growth of the primary tumor, we observed that targeting glutamine metabolism significantly reduced lung metastasis (Figure 6A-C). The ability of JHU083 to inhibit metastasis was also observed when we delivered tumor via tail vein injection (Supplementary Figure 5A). Since MDSCs are believed to play a crucial role in facilitating metastasis, we interrogated the lungs of JHU083 treated and untreated mice for MDSCs. We observed an increase in the CD8:MDSCs ratios in the lungs of the treated versus untreated mice (Figure 6D). In addition to FACS, we also performed metabolomics on the lungs from the JHU083 treated and untreated mice on day 17, <u>before</u> visible metastasis occurs to understand the possible metabolic changes related to development of metastasis. Similar to the primary tumors, LC-MS analysis of the lungs revealed two distinct metabolic clusters (Figure 6E and F). That is, despite a <u>lack of macroscopic metastasis</u> in the lungs on day 17, we observed significant metabolic changes (Figure 6F and Supplementary Figure 5B). Strikingly, in agreement with our primary tumor data, the kynurenine level was markedly reduced in the lungs from the JHU083-treated mice (Figure 6G). Interestingly, we observed higher IDO expression in the lungs from untreated mice with tumors prior to the appearance of metastases compared to the lungs of tumor-free mice. Similar to primary tumor, JHU083 treatment decreased IDO expression in the lung (Figure 6H). Overall, these data suggest that the robust ability of targeting glutamine metabolism to inhibit metastasis may be attributed in part to altering the “metabolic” and immunologic metastatic microenvironment.

**Figure 6.**
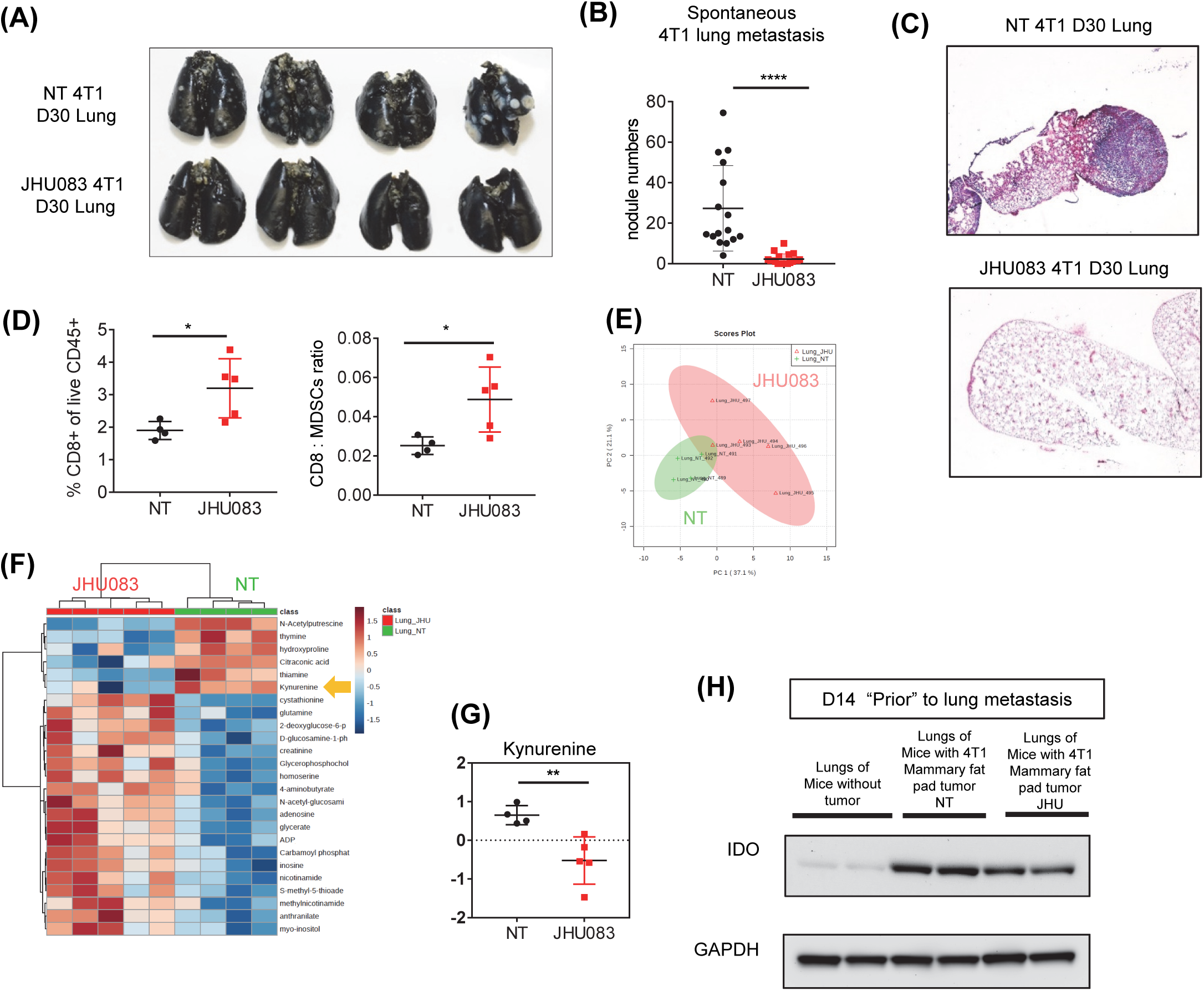
Reduced spontaneous lung metastasis along with metabolic changes of lungs from vehicle vs. JHU083 treated 4T1 tumor-bearing mice. 0.1×10^6^ 4T1 cells were implanted subcutaneously into mammary fat pad of BALB/cJ female mice. The whole lung were harvested, and spontaneous lung metastasis were analyzed (A-D). (A-B) To quantify tumor nodules, On day 30, lungs were inflated with 15% india ink. (A) Representative picture of lungs. (B) Quantification of tumor nodules in lungs. (C) Representative histology sections stained with H&E. (D) Percentages of CD8^+^ of live CD45^+^ cells in lung were analyzed by flow cytometry (left). Ratio of CD8 cells to MDSCs in 4T1 tumor (right). (E-H) On day 17, the whole lungs were harvested, and (E-G) whole lung lysates were applied for LC-MS analysis. (E) Principal component analysis between the vehicle treated (NT) (green) and JHU083 (red) and (F) Heatmap visualization of the metabolite changes between NT (green) and JHU083 (red) treated group were shown. (G) Relative amounts of kynurenine between NT and JHU083 treated group. (H) On day 14, IDO expression on lung lysates from tumor free, and 4T1 tumor-bearing mice with or without JHU083 treatment. Data are from three independent experiment with 4-6 mice per group (A,C,D and H), combined three independent experiments with 4-6 mice per group (B) or from one experiment with 4-5 mice per group (E-G). *P < 0.05, **P < 0.01, ****P<0.001, MEAN ± S.D. t-test (G) and Mann-Whitney t tests (B and D).

### The glutamine antagonist JHU083 enhances the efficacy of checkpoint blockade

Our studies demonstrate that targeting glutamine metabolism boosts immune responses by reprogramming tumor metabolism, enhancing a pro-inflammatory phenotype of TAMs, reducing MDSCs, and promoting ICD. Recall, 4T1 tumor cells are extremely resistant to immunotherapy in the form of checkpoint blockade (Figure 1A). Thus, we were interested in determining if blocking tumor glutamine metabolism could enhance the efficacy of checkpoint blockade in this model. First, we tested this hypothesis on a checkpoint blockade sensitive tumor. We assessed the ability of JHU083 to inhibit growth of EO771, which is similar to 4T1 triple negative breast cancer. The EO771 tumor model is moderately sensitive to immunotherapy in the form of anti-PD1 monotherapy or anti-CSF1R + anti-CD40 combination immunotherapy (Hoves et al., 2018). We observed significant inhibition of tumor growth and enhanced survival with JHU083 treatment alone (Supplementary figure 6A and B). Furthermore, anti-PD1 monotherapy resulted in delayed tumor growth. Notably, the combination of JHU083 + anti-PD1 or JHU083 + anti-PD1 + anti-CTLA4 resulted in greater inhibition of tumor growth and enhanced survival suggesting an additive or synergistic effect of combining metabolic therapy with checkpoint blockade (Supplementary figure 6A and B).

This result prompted us to evaluate whether JHU083 can enhance the efficacy of checkpoint blockade even in tumors that are resistant to immunotherapy. To this end, mice were injected with 4T1 tumors and treated on day 7 post injection with either vehicle, JHU083 alone, anti-PD1+ anti-CTLA4, or JHU083 + anti-PD1 + anti-CTLA4. The mice treated with anti-CTLA4 and anti-PD1 had no therapeutic benefit compared to the vehicle treated group as seen in Figure 1 (Figure 7A). The JHU083-treated group displayed delayed tumor growth and an increase in survival (Figure 7A and B). Strikingly, when mice were treated with JHU083, anti-PD1 and anti-CTLA4, we observed further attenuation of tumor growth and an increase in survival compared to the JHU083 monotherapy group (Figure 7A and B). Taken together, these findings demonstrate that by altering the TME glutamine inhibition can enhance the efficacy of checkpoint blockade even in tumors that are resistant to immunotherapy (Figure 7C).

**Figure 7.**
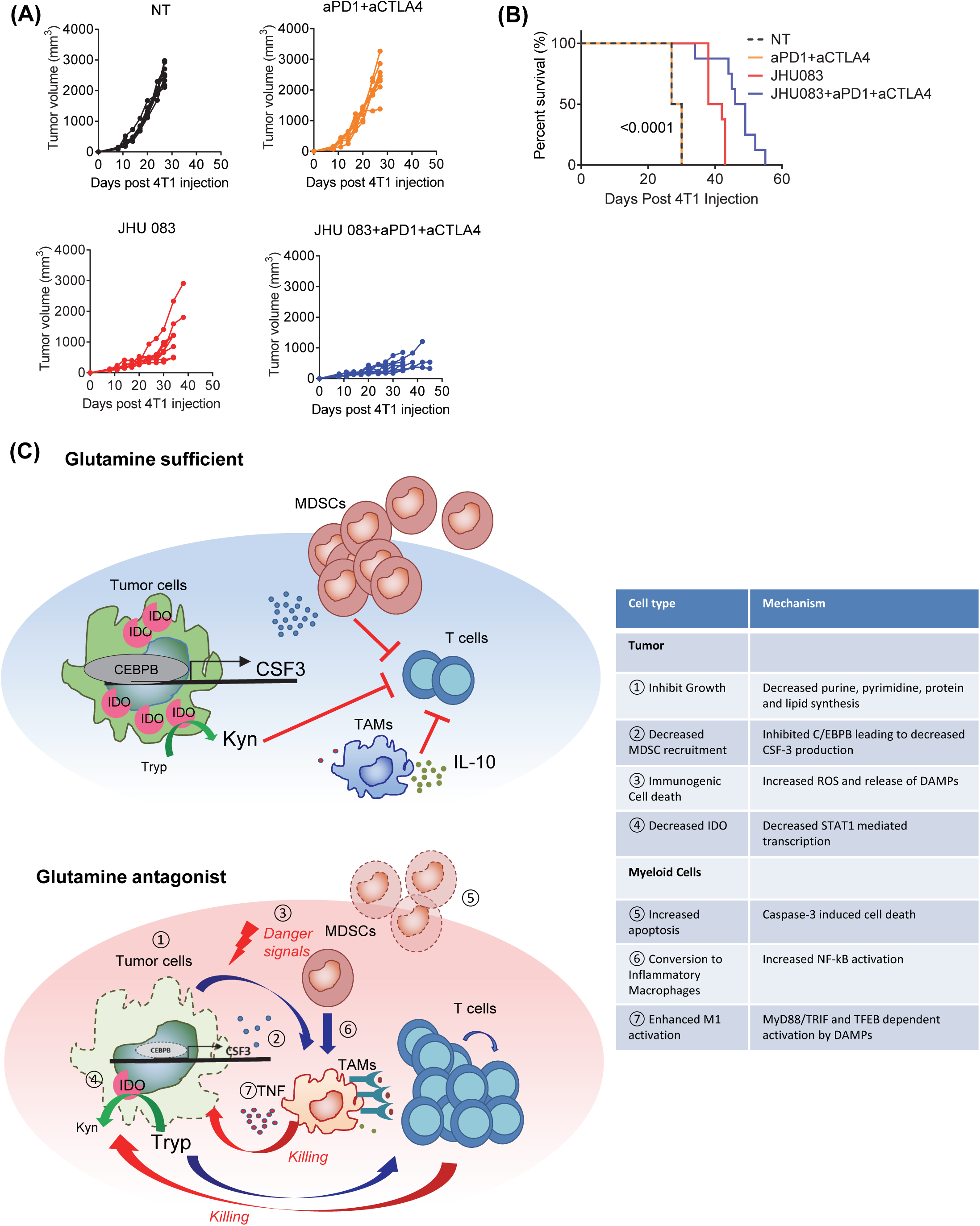
Glutamine antagonism enhances immunotherapy in 4T1 tumor-bearing mice. (A-B) 4T1 tumor-bearing mice were treated with JHU083 alone or JHU083 in combination with 250 μg anti-PD1 and 100 μg of anti-CTLA4 antibodies (On day 7, 10, 13, 17, and 24). (A) Tumor sizes and (B) survival curves were recorded. Data are representative of three independent experiments. N=8 mice per group (C) Proposed model. MEAN ± S.D. Log-rank (Mantel-Cox) test (B).

## Discussion

It is becoming increasingly clear that specialized tumor metabolism is not merely to support the growth and energetics of cancer cells but also plays a critical role in creating an immunosuppressive TME. To this end, the metabolic program of suppressive myeloid cells is specialized in order to thrive within the TME. We hypothesize that targeting glutamine metabolism would not only inhibit tumor growth but also alter the TME and subsequently tumor immune evasion. Therefore, targeting glutamine metabolism led to **i.** inhibition of tumor derived CSF-3 **ii.** Inhibition of tumor IDO expression resulting in decreased kynurenine **iii.** Immunogenic cell death of tumor cells **iv.** Apoptosis of MDSC **v.** Conversion of MDSC to inflammatory macrophages **vi**. Enhanced activation of macrophages and antigen presentation. The net result of these effects was decreased tumor growth and enhanced anti-tumor immunity. Importantly, our studies reveal the intimate relationship between the metabolism of tumor cells and the metabolism of suppressive immune cells and how targeting glutamine metabolism can alleviate immune evasion.

The critical role of glutamine in supporting the prodigious anabolic requirements of cancer cells has been appreciated for some time (Altman et al., 2016; Galluzzi et al., 2013; Tennant et al., 2010). Current efforts in targeting glutamine in tumors has primarily focused on the initial step of glutaminolysis through the development of selective GLS inhibitors (Gross et al., 2014; Wang et al., 2010; Xiang et al., 2015). While such inhibitors demonstrate robust efficacy in vitro, it is becoming clear that glutaminase-targeted therapy is far less effective in vivo (Biancur et al., 2017; Davidson et al., 2016; Gao et al., 2009; Romero et al., 2017). As such, we have developed a novel pro-drug of the glutamine antagonist DON which not only inhibits glutaminase but also all other glutamine requiring reactions important to tumor growth including purine and pyrimidine biosynthesis, redox control, glycosylation, amino acid and collagen synthesis, autophagy and epigenetics (Altman et al., 2016; Yang et al., 2017). DON as an anti-tumor agent has been studied for 60 years. While DON treatment resulted in some encouraging responses in phase I and II clinical trials in the 1950s to the 1980s, the development of DON was limited by its GI toxicity (Ahluwalia et al., 1990; Lemberg et al., 2018). Our novel compound, JHU083, limits toxicity by creating an inert prodrug that is preferentially (though not exclusively) converted to the active compound DON within the TME (Lemberg et al., 2018; Rais et al., 2016). It is important to note, that while we can evaluate the efficacy of our approach in mice, we cannot evaluate the toxicity and pharmacokinetics in small animals (rodents) because they metabolize the prodrugs differently than humans (Rais et al., 2016). As such, our dosing schedule in mice is much more limited by the potential toxicities than it would be in humans. Nonetheless, even though JHU083 is rapidly converted to DON in mice, we have identified a robust therapeutic window to evaluate its effects on tumor growth and the TME. Furthermore, unlike the previous clinical trials employing DON, in the modern era, our studies provide robust clinical rationale for developing combination regimens using our novel DON prodrug along with immunotherapy.

While the specialized metabolism of tumors promotes growth it also profoundly influences the TME. Indeed, the hypoxic, acidic, nutrient depleted TME in and of itself serves to inhibit anti-tumor immune responses. Such an environment favors the residency of suppressive myeloid cells such as MDSCs, TANs and TAMs all of which contribute to promoting tumor growth, angiogenesis, metastasis, and immune escape (Binnewies et al., 2018). Additionally, suppressive myeloid cells contribute to resistance against immune checkpoint blockade (Sharma et al., 2017; Steinberg et al., 2017). Our data demonstrate that targeting glutamine metabolism leads to a marked decrease in MDSCs in both the peripheral blood of tumor bearing mice and within the tumors itself. Mechanistically this is due in part to 1) increased caspase3 dependent cell death, 2) the decreased secretion of CSF-3 from the both tumors and TAMs by reduced transcription factor C/EBPB via increased autophagy dependent degradation, and 3) MDSCs differentiation into pro-inflammatory TAMs. Interestingly, glutamine antagonism did not simply reduce the percentage and absolute numbers of TAMs within the tumor. Rather, it promoted the generation of pro-inflammatory TAMs. Analogous with our findings are recent studies that demonstrate that glutamine depletion enhances M1 and reduces M2 macrophage phenotype and function (Liu et al., 2017).

A recent study demonstrated that inhibition of aerobic glycolysis in the tumor can reduce MDSC recruitment by reduction of CSF3 secretion (Li et al., 2018). However, directly targeting glycolysis will also inhibit pro-inflammatory (M1) macrophage function. On the other hand, by targeting glutamine metabolism we can achieve the same goal with regard to inhibiting MDSC but simultaneously promoting the generation of pro-inflammatory macrophages and thus enhancing anti-tumor immunity. Thus, our paper provides valuable insight regarding the translational impact of targeting glutamine metabolism from both perspectives to the dynamic interplay between the tumor and also the immune cells.

In as much as glutamine plays a critical role in multiple anabolic metabolic pathways, we hypothesized that glutamine antagonism would drastically alter the “metabolic” TME. Indeed, in 4T1 tumors from the JHU083 treated mice, we observed decreases of: citrulline, N-carbamoyl-L-aspartate, thymine, S-adenosyl-L-methionine, homoserine, guanosine, nicotinamide ribotide, hydroxyproline and succinate. Surprisingly, of the 200 metabolites queried, kynurenine was the most differentially regulated. Kynurenine is the byproduct of tryptophan metabolism by IDO and has potent immunosuppressive effects. IDO knockout mice robustly reject tumors and inhibitors of IDO are being developed clinically as immunotherapy (Holmgaard et al., 2013; Munn and Mellor, 2016; Uyttenhove et al., 2003). Unexpectedly, JHU083 inhibited conversion of tryptophan to kynurenine. However, its mechanism of action was not by directly inhibiting IDO but rather by inducing the down modulation of IDO transcriptions via reduced STAT1 and STAT3 transcriptional activity. While the pathways that lead to this inhibition are not precisely clear, glutamine metabolism is critical for a number of processes important in post-translational modification including hexosamine biosynthesis, purine and pyrimidine biosynthesis, redox control, and amino acid synthesis. The disruption of these processes through glutamine blockade likely has significant effects on the transcriptional activity of STAT1 and STAT3, which are also known to be highly regulated by a wide range of post-translational modifications.

In addition to inhibiting growth of the primary tumor, glutamine antagonism proved to be a potent means of inhibiting the development of metastasis. This observation has important clinical relevance to many tumors (especially breast cancer) where metastatic spread of the primary tumor negates successful surgical removal. In the 4T1 model, a major site of metastasis is the lung. Interestingly, we observed both metabolic and immunologic differences in the lungs of untreated and treated mice even in the absence of macroscopic metastasis. MDSCs are thought to play an integral role in promoting metastasis (Condamine et al., 2015; Huang et al., 2013; Yu et al., 2013). It has been shown that MDSCs increase angiogenesis, tumor invasion, and formation of a pre-metastatic niche by enhancing pro-angiogenetic factor (such as VEGF, PDGF, b-FGF, and angiopoietins), and MMPs, chemokines (such as CXCL1, CXCL2, MCP1 and CXCL5) (Condamine et al., 2015). To this end, we observed an increase in the CD8:MDSCs ratio in the lungs of the treated mice even in the absence of observable tumor. Likewise, kynurenine levels were decreased in the lungs of JHU083 treated mice compared to untreated mice even before there was evidence of macroscopic metastasis. Previous studies have shown that kynurenine can promote metastasis by inducing epithelial-to-mesenchymal transition by activating the aryl hydrocarbon receptor (Xue et al., 2018).

Immunotherapy in the form of anti-PD1 and anti-CTLA4 has revolutionized our approach to treat certain cancers. Yet, in spite of these successes it is clear that not all cancers respond to checkpoint blockade alone and even amongst responsive cancers, not all patients respond (Del Paggio, 2018; Larkin et al., 2015; Sharma et al., 2017; Tumeh et al., 2014). Such observations point to multiple different mechanisms of tumor immune evasion. Our data suggest that by targeting glutamine metabolism we can enhance the efficacy of immunotherapy. To this end, in the anti-PD1 responsive EO771 model, the addition of JHU083 markedly enhanced the overall response rate of checkpoint blockade. Furthermore, in the 4T1 model that was resistant to combined anti-PD1 and anti-CTLA4 treatment, we could overcome resistance in part by blocking glutamine metabolism. Overall, these observations support the view that tumor metabolism represents a means by which cancer cells can evade anti-tumor immune responses. Further, we provide the preclinical rationale for strategies involving targeting glutamine metabolism as a means of enhancing immunotherapy for cancer.

## Experimental Procedures

### Animal

C57BL/6, CD45.1 BALB/cJ, OTI, RAG1 knock out and BALB/cJ (both male and female mice, 6-8 weeks of age) were purchased from Jackson Laboratories. Mice were randomly assigned to experimental groups. NSG mice were obtained from Johns Hopkins Animal resources facility. MyD88/TRIF double knock out mice were kindly provided by Dr. Franck Housseau (Johns Hopkins University) (Hoebe et al., 2003; Kawai et al., 1999). TFEB knock out mice were kindly provided by Dr. Andrea Ballabio (Baylor College of Medicine) (Settembre et al., 2012). The Institutional Animal Care and Use Committee of Johns Hopkins University (Baltimore, MD) approved all animal protocols.

### Chemical compound

6-diazo-5-oxo-l-norleucine (DON) was purchased from Sigma-Aldrich. JHU083 (Ethyl 2-(2-Amino-4-methylpentanamido)-DON) was synthesized using our previously described method. (Nedelcovych et al., 2017; Rais et al., 2016).

### Cell lines and tumor model

4T1 breast cancer cell lines, 3LL lung carcinoma cell lines, and RAW 264.7 macrophages cell lines were purchased from the ATCC. EO771 breast cancer cell lines were purchased from CH3 BioSystems. MC38 OVA, B16 OVA cell lines were kindly provided by Dr. Drew Pardoll (Johns Hopkins University). 4T1 cells and EO771 cells were cultured in RPMI supplemented with 10% FBS, 1% penicillin/streptomycin, and 10 mM HEPES and 3LL, RAW 264.7 and MC38 OVA cells were cultured in DMEM supplemented with 10% FBS, 2 mM glutamine, 1% penicillin/streptomycin, and 10 mM HEPES. All cell lines were regularly tested to confirm mycoplasma free using MycoAlert mycoplasma detection kit (Lonza). Cells were never passaged more than 3 weeks before use in an experiment. 4T1 cells (1 × 10^5^ cells in 200 μl per mouse) were subcutaneously inoculated into the mammary fat pad of BALB/cJ mice. EO771 cells (2 × 10^5^ cells in 200 μl per mouse) were subcutaneously inoculated into the mammary fat pad of C57BL/6 mice. 3LL (5 × 10^5^ cells in 200 μl per mouse) cells were subcutaneously inoculated into right flank of C57BL/6 mice. Tumor-bearing mice were treated with vehicle (PBS, referred as a NT group) or glutamine antagonist prodrug, JHU083 (1mg/kg), daily starting at day 7 or day 9 after tumor inoculation. After 7 days of 1mg/kg treatment, a lower dose (0.3 mg/kg) of JHU083 was used. On day 7, 10, 13, 17, and 24, 4T1 tumor-bearing mice were injected intraperitoneally with 250 μg anti-PD1 (InVivoPlus RMP1-14, BioXCell) and/or 100μg anti-CTLA4 antibodies (InVivoPlus 9H10, BioXCell). Tumor burdens were monitored every 2-4 days by measuring length and width of tumor. Tumor volume was calculated using the formula for caliper measurements: tumor volume = (L × W^2^)/2, where L is tumor length and is the longer of the 2 measurements and W is tumor width. Mice were euthanized when tumor size exceeded 2 cm in any dimension or when the mice shows the hunched posture, ruffled coat, neurological symptoms, severe weight loss, labored breathing, weakness or pain.

To establish a pulmonary metastasis model, 4T1 cells (1 × 10^5^ cells in 200 μl per mouse) were injected into the BALB/cJ via tail vein. Tumor-bearing mice were treated with vehicle or JHU083 (1mg/kg) starting at day 2 after tumor inoculation. After 7 days of 1mg/kg treatment, lower dose (0.3 mg/kg) of JHU083 was used.

To assess MDSC conversion into TAMs, MDSCs from the blood were isolated from CD45.1 4T1 tumor bearing mice (21 days after 4T1 tumor inoculation). Isolated MDSCs were adoptively transferred into CD45.2 4T1 tumor bearing mice (7days after 4T1 tumor inoculation). Then, recipient CD45.2 4T1 tumor bearing mice were treated with JHU083 (1mg/kg) until harvesting tumors on day 7.

To track tumor cells in vivo, the green fluorescent protein (GFP) expressing 4T1 tumor cells were generated by transducing cells with lentiviral vector carrying GFP gene (phage ubc nls ha pcp gfp plasmid, Addgene Plasmid #64539).

### Tumor from primary tumor and spontaneous pulmonary metastasis digestion and sorting

Tumors or lungs from tumor bearing mice were minced in RPMI with 2% FBS, 2 mg/mL collagenase IV (GIBCO) and DNase (Roche), and incubated in 37°C for the digestion. After 30 minutes, 0.02% EDTA was added to inhibit further digestion, and cells were filtered through 70 µm cell strainer to remove debris. For flow cytometry, single-cell suspensions were washed with PBS, and incubated in ammonium-chloride-potassium lysis buffer (Quality Biological). For sorting of TAMs, cell pellet was suspended in 30% Percoll and then carefully layered onto 70% Percoll. Samples were centrifuged at 2000 rpm at room temperature without brake for 30 minutes. After washing, tumor-infiltrating cells were isolated using CD45 isolation kit (Miltenyl). After staining, tumor associate macrophages (CD45^+^ 7AAD^-^ CD8^-^ Ly6C^-^ Ly6G^-^ CD11b^+^ F4/80^+^) were sorted using BD FACSAria™ Fusion.

### Flow cytometry

Single cell-suspensions were stained with antibodies after Fc blocking (BD bioscience). The following antibodies and staining reagents were purchased from Biolegend: anti-CD45 (30-F11), anti-F4/80 (BM8), anti-CD11b (M1/70), Ly6C (HK1.4), Ly6G (RB6-8C5), MHC Class II (M5/114.15.2), CD8 (53-6.7), CD4 (RM4-5), Cell signaling: Calreticulin (D3E6), Thermofisher: iNOS(CXNFT), LIVE/DEAD® Fixable Near-IR Dead Cell Stain Kit, TLR4 (UT41), CellROX™ Deep Red Flow Cytometry Assay Kit, Fixation and Permeabilization Buffer Set, and BD Bioscience: TNF (MP6-XT22), GM-CSF(MP1-22E9), 7-AAD, BD, BD Cytofix/Cytoperm Plus Kit (with BD GolgiPlug) and staining were followed manufacture’s protocol. Cells were acquired using BD FACSCalibur or BD FACSCelesta, and data were analyzed using FlowJo (FlowJo, LLC).

### RNA sequencing and data analysis

On day 14 after tumor inoculation, TAM (CD45^+^ 7AAD^-^ CD8^-^ Ly6C^-^ Ly6G^-^ CD11b^+^ F4/80^+^) from vehicle or JHU083 treated 4T1 tumor-bearing mice were sorted on BD FACSAria™ Fusion. For RNA sequencing analysis, total RNA (5 mice per group) was extracted using RNeasy Micro Kit (QIAGEN). Samples were sent to Admera Health for sequencing and analysis. Poly(A)^+^ transcripts were isolated by NEBNext® Poly(A) mRNA Magnetic Isolation kit. Prior to first strand synthesis, samples are randomly primed and fragmented using NEBNext® Ultra™ RNA Library Prep Kit for Illumina®. The first strand was synthesized with the Protoscript II Reverse Transcriptase. Samples were pooled and sequenced on a HiSeq with a read length configuration of 150 PE. The transcriptomic analysis work flow began with a thorough quality check by FastQC v0.11.2. The latest reference genome (GRCM38) were used for read mapping. The statistical significant gene analysis in context of gene Ontology and other biological signatures were performed using Gene Set Enrichment Analysis (GSEA) and DAVID. The RNA sequencing data have been deposited in the GEO under ID codes GSE119733.

### The Cancer Genome Atlas data analysis

To determine the correlation between IDO and glutamine utilizing enzymes which is inhibited by DON expression levels, the cancer atlans data (TCGA) were used. Breast invasive carcinoma (N=1085) and normal (N=112) samples were analyzed by using GEPIA (Gene Expression Profiling Interactive Analysis, http://gepia2.cancer-pku.cn/=index) (Tang et al., 2017).

### Generation of BMDMs

For preparation of bone marrow cell suspensions, the bones of both hind limbs (two tibias and two femurs) were flushed with ice-cold DMEM supplemented with 10% FBS, 1% penicillin/streptomycin and 2 mM L-glutamine (cell media) plus 20% L929-conditioned media. The cells were incubated at 37 °C, and on day 4, non-adherent cells were removed, and replaced with the fresh L929 conditioned media. On day 7, BMDMs were lifted using Cellstripper (Mediatech, Manassas, VA). 1 × 10^6^ cells BMDMs were seeded in 12-well plates, and treated with DON. To make tumor-conditioned media, tumor cells were cultured in the presence or absence of DON (0.5 μM or 1 μM). After 1 hour of incubation, cells were washed and replaced with drug-free fresh media. After 24 hours, supernatants were harvested and used as conditioned media (CM). BMDMs were cultured in the presence of these conditioned media for 24 hours.

### T cell priming assay by co-culture with OTI and tumor cells

3 × 10^5^ BMDM cells were plated on a 12 well plate, then B16 OVA or MC38 OVA 50,000 cells were added. After attachment, different doses of DON were applied and incubated for 24 hours. CD8^+^ from OTI mice were isolated by CD8^+^ T Cell Isolation Kit (Miltenyi Biotech). Purified OTI CD8^+^cells or whole splenocytes from OT1 mice were labeled with Cell Proliferation Dye eFluor™ 450 (Invitrogen). After discarding the supernatant, labeled 3 × 10^5^ CD8^+^ or 2 × 10^6^ whole splenocytes were co-cultured with BMDMs and tumor cells. After 72 hours, cells were analyzed by flow cytometry.

### Immunoblotting

For immunoblotting of MDSCs, MDSCs from the blood were isolated from 4T1 tumor bearing mice (more than 21 days after 4T1 tumor inoculation. For the nuclear and cytoplasm fractionations, 10 × 10^6^ BMDMs were washed twice with PBS, then lysed cells in cytoplasm separation buffer composed of 10 mM HEPES, 60 mM KCl, 1 mM EDTA, 0.075% (v/v) NP40, 1 mM DTT and 1 mM PMSF for cytoplasm fraction, and RIPA buffer with NaF, protease inhibitor, PMSF, sodium pyrophosphate, beta glycerophosphate and sodium vanadate for nuclear fraction. For immunblotting, cells were lysed in RIPA buffer with NaF, protease inhibitor, PMSF, sodium pyrophosphate, beta glycerophosphate and sodium vanadate. Western bloting was performed using a standard protocol (Life Technologies). The following antibodies were used from Cell Signaling: anti-active caspase 3, anti-p-STAT3 (Tyr705), anti-STAT3, anti-p-NF-κB p65 (Ser536), anti-C/EBPB, anti-ULK (Ser 757), anti-ULK (Ser 555), anti-ULK1, anti-p-AMPK (Thr172), anti-p-S6K (Thr389), anti-LC3, anti-Histon H3, anti-IDO, anti-LaminB, and anti-β-actin, and from Abcam: anti-LAMP2 (GL2A7). All images were captured and analyzed using UVP BioSpectrum 500 Imaging System.

### Enzyme-linked immunosorbent assay

Serum or cell culture supernatants were collected, and CSF3, TNF, and IL-10 were analyzed by ELISA as described by the manufacturer (ThermoFisher).

### Metabolite Extractions

Tumor (95 mg) and lung (whole lung) were harvested on day 17 from 4T1 tumor bearing mice treated with or without JHU083. To minimize ischemic time, the organs were harvested within 1 minute. Metabolites were extracted from tumor and lung in a methanol:water (80:20, v/v) extraction solution after homogenization with an ultrasonic processor (UP200St, Hielscher Ultrasound Technology). Samples were vortexed and stored at −80℃ for at least 2 hours to precipitate the proteins. The metabolite containing supernatant was isolated after centrifugation at 15,000g for 10 minutes and dried under nitrogen gas for subsequent analysis by LC-MS.

### Metabolite Measurement with LC-MS

Targeted metabolite analysis was performed with LC-MS as previously described. Dried samples were re-suspended in 50% (v/v) acetonitrile solution and 4μL of each sample were injected and analyzed on a 5500 QTRAP triple quadrupole mass spectrometer (AB Sciex) coupled to a Prominence ultra-fast liquid chromatography (UFLC) system (Shimadzu). The instrument was operated in selected reaction monitoring (SRM) with positive and negative ion-switching mode as described. This targeted metabolomics method allows for analysis of over two hundreds of metabolites from a single 25min LC-MS acquisition with a 3ms dwell time and these analyzed metabolites cover all major metabolic pathways. The optimized MS parameters were: ESI voltage was +5,000V in positive ion mode and –4,500V in negative ion mode; dwell time was 3ms per SRM transition and the total cycle time was 1.57 seconds. Hydrophilic interaction chromatography (HILIC) separations were performed on a Shimadzu UFLC system using an amide column (Waters XBridge BEH Amide, 2.1 × 150 mm, 2.5μm). The LC parameters were as follows: column temperature, 40 ℃; flow rate, 0.30 ml/min. Solvent A, Water with 0.1% formic acid; Solvent B, Acetonitrile with 0.1% formic acid; A non-linear gradient from 99% B to 45% B in 25 minutes with 5min of post-run time. Peak integration for each targeted metabolite in SRM transition was processed with MultiQuant software (v2.1, AB Sciex). The preprocessed data with integrated peak areas were exported from MultiQuant and re-imported into Metaboanalyst software for further data analysis (e.g. statistical analysis, fold change, principle components analysis, etc.).

### Immunohistochemistry

Lungs were inflated with 10% neutral buffered formalin by tracheal cannulation. Lungs were excised and placed in formalin for 24 hours. Formalin fixed lung section were paraffin-embedded and processed for histological analysis. Lung section were stained with hematoxylin and eosin (H&E), and images were captured and analyzed by microscope at 40 × magnification.

### Lung metastasis analysis

Lung metastases were analyzed by inflation with 15% india ink. After intra-tracheal injection of india ink, lungs were harvested and washed with in Feket’s solution (70% ethanol, 3.7% paraformaldehyde, 0.75 M glacial acetic acid). Lungs were placed in fresh Feket’s solution overnight, and surface white tumor nodules were counted in a group-blinded fashion using a Nikon stereomicroscope.

### Statistics

Generating graphs and statistical analysis were performed with Prism 7 (GraphPad). Comparison between two means was done by *t*-test or non-parametric 2-tailed Mann-Whitney *t*-test. Comparisons between three or more means were done by Kruskal-Wallis test with Dunn’s multiple comparisons post-test. Survival test was done by Log-rank (Mantel-Cox) test. The association between two ranked variables was done by spearman rank correlation.

## Author Contributions

M.O., I.S., R.L., I.S., W.X., S.L.C., A.J.T., R.L.B., C.H.P., J.E., M.L.A., J.W. and Y.C. performed and analyzed experiments. L.Z. performed and analyzed LC-MS/MS experiments. P.M., R.R., B.S.S designed and synthesized JHU083. M.O. and J.D.P wrote the manuscript. M.R.H. and J.D.P. supervised the project.

## Acknowledgements

We thank the members of the Horton and Powell labs for review of this manuscript. We thank Lee Blosser for assistance with flow cytometry sorting. This work was supported by the NIH grant (90079285 R01 to J.D.P. and B.S.S. and S10 OD016374 to the JHU Microscopy Facility), the Bloomberg∼Kimmel Institute for Cancer Immunotherapy to J.D.P. and B.S.S., Under Armour Women’s Health & Breast Cancer Innovation Grants to J.D.P..

---

**FigS1 related to Figure 1.**
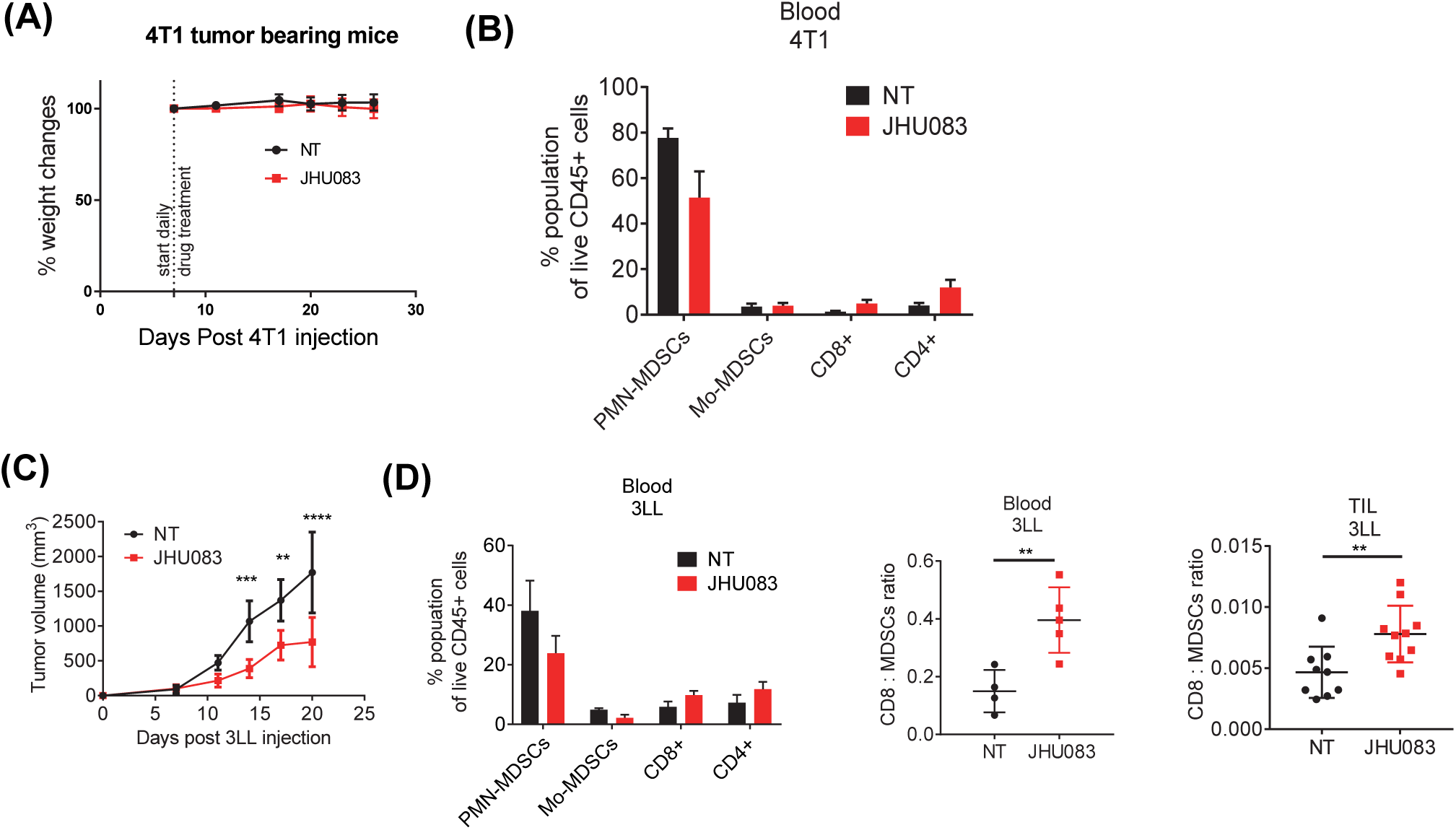
Glutamine antagonist inhibits 3LL tumor growth by enhancing CD8+ population, and reducing PMN-MDSCs and Mo-MDSCs. 0.1×10^6^ 4T1 cells were implanted subcutaneously into the mammary fat pad in BALB/cJ female mice. 4T1 tumor-bearing mice were treated with JHU083 (1mg/kg) starting at day 7 after tumor inoculation. After 7 days of treatment, a lower dose (0.3 mg/kg) of JHU083 was used (A-B). (A) Mice weights were monitored and recorded. (B) Percentages of PMN-MDSCs, Mo-MDSCs, CD8+, and CD4+ of live cells from blood in 4T1 tumor-bearing mice were analyzed by flow cytometry (N=9-10/group). 0.5×10^6^ 3LL cells implanted subcutaneously into the right flank of C57BL/6J male mice. 3LL tumor-bearing mice were treated with JHU083 (1mg/kg) starting at day 7 after tumor inoculation. After 7 days of treatment, a lower dose (0.3 mg/kg) of JHU083 was used (C-D). (C) 3LL tumor sizes were measured. (D) Percentages of PMN-MDSCs, Mo-MDSCs, CD8^+^, and CD4^+^ of live cells from blood in 3LL tumor bearing mice were analyzed by flow cytometry at day 15. Ratio of CD8^+^ cells to MDSCs from blood and TIL. **P < 0.01, ***P < 0.005 Mann-Whitney t tests.

**FigS2 related to Figure 3.**
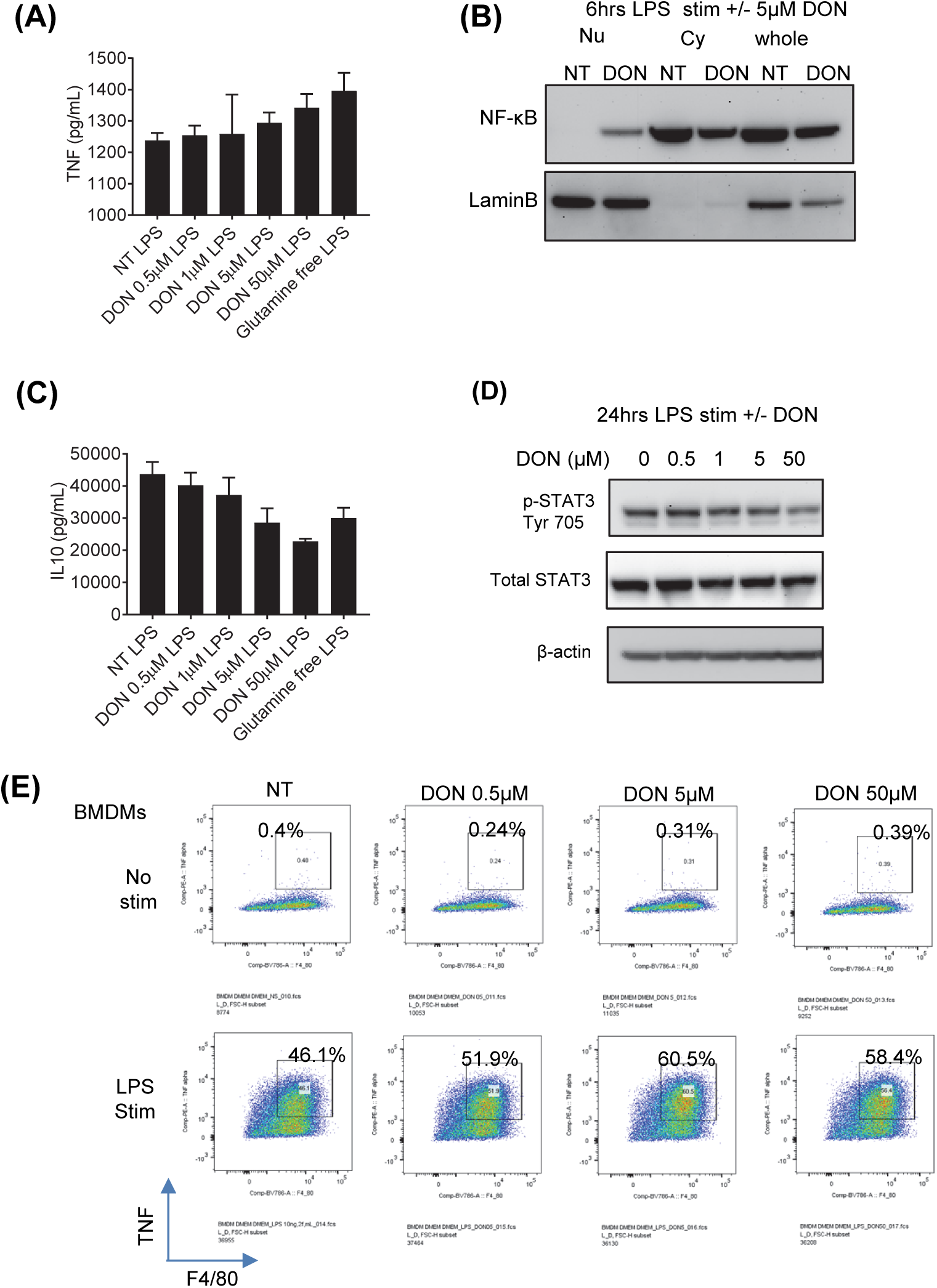
Glutamine antagonist treated BMDMs enhance TNF but reduce IL-10 secretion. BMDMs were stimulated with LPS in the presence or absence of indicated dose of DON (glutamine antagonist) or in glutamine free media (GF) for 6 or 24 hours. (A) After 24 hours, supernatants were collected and TNF levels were measured by ELISA. (B) BMDM were stimulated with LPS and treated with or without DON. NF-κB levels were probed in nuclear (Nu) and cytosolic(Cy) fraction from LPS or LPS+5 μM DON treated (6 hours) BMDMs by immunoblotting. LaminB was used to confirm the nuclear fraction. (C) IL-10 in supernatants from (A) were measured by ELISA. (D) p-STAT3 (Tyr705) level and total STAT3 from BMDMs stimulated with either LPS, LPS+DON, or LPS in glutamine free media (GF) for 24 hours were measured by immunoblotting. (E) BMDMs were stimulated with LPS and Golgi-plug in the presence or absence of indicated dose of DON for 9 hours. TNF levels were measure by flow cytometry.

**FigS3 related to Figure 4.**
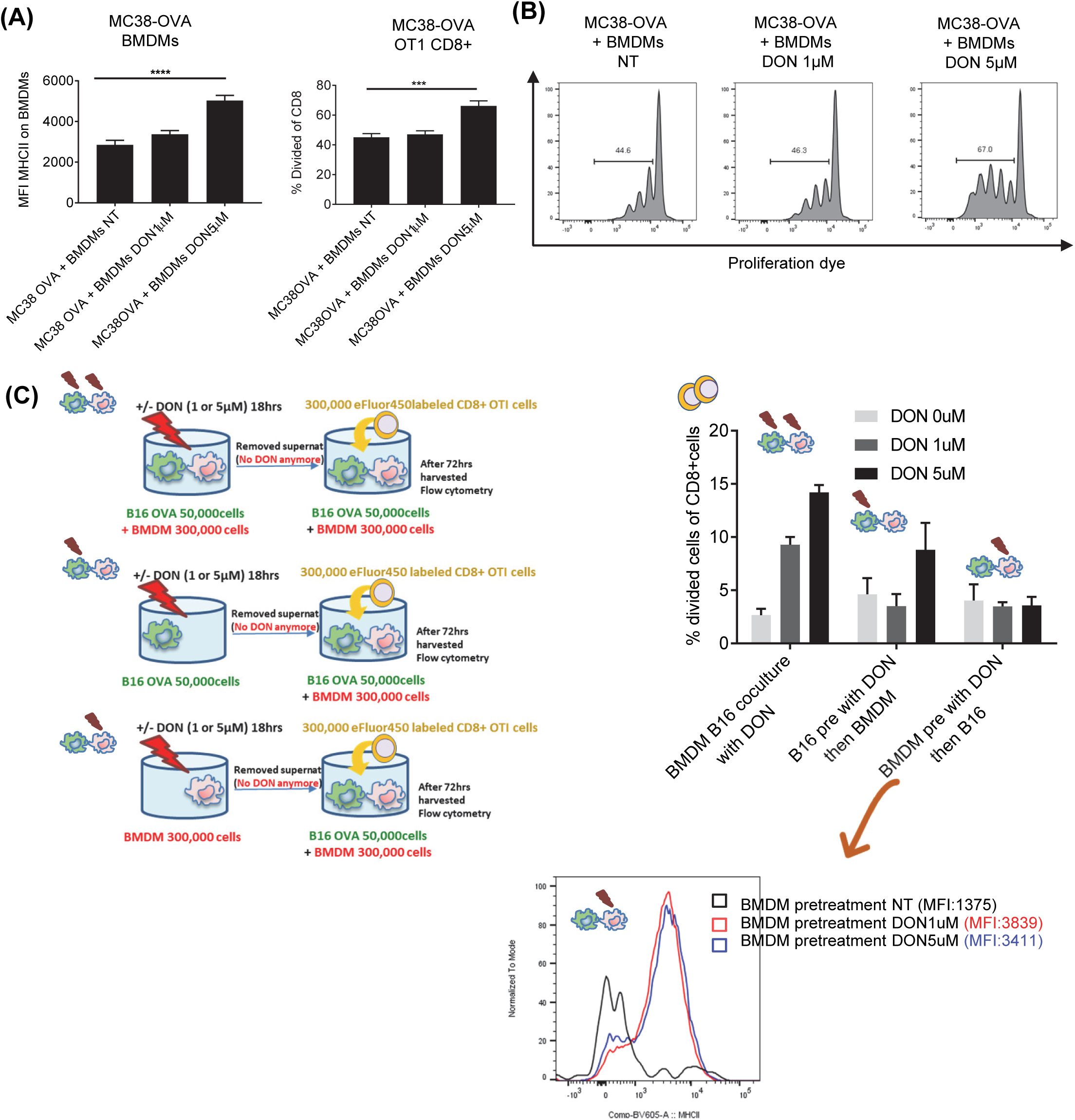
Glutamine antagonist treated BMDMs enhance antigen presentation. (A and B) 0.3×10^6^ BMDMs and 5×10^4^ MC38-OVA tumor cells were co-cultured in the presence or absence of 1 μM or 5 μM of DON. After 24 hours of incubation, supernatants were discarded and 2×10^6^ eFluor450-labeled whole splenocytes from OTI mice were added. (A) MHCII expression on BMDMs and percentages of divided cells from CD8+ population were analyzed by flow cytometry. (B) Histogram showing T cell proliferation (C) BMDMs or B16-OVA tumor cells were cultured in the presence or absence of 1 μM or 5 μM of DON. After 18 hours of incubation, 0.3×10^6^ BMDMs or 5×104 B16-OVA tumor cells were seeded and 0.3×106 eFluor450-labeled CD8+ from OTI mice were added. Schematic of the experiment (Left) Percentages of divided cells from CD8+ population were analyzed by flow cytometry (right). Histogram showing MHCII expression (bottom)

**FigS4 related to Figure 5.**
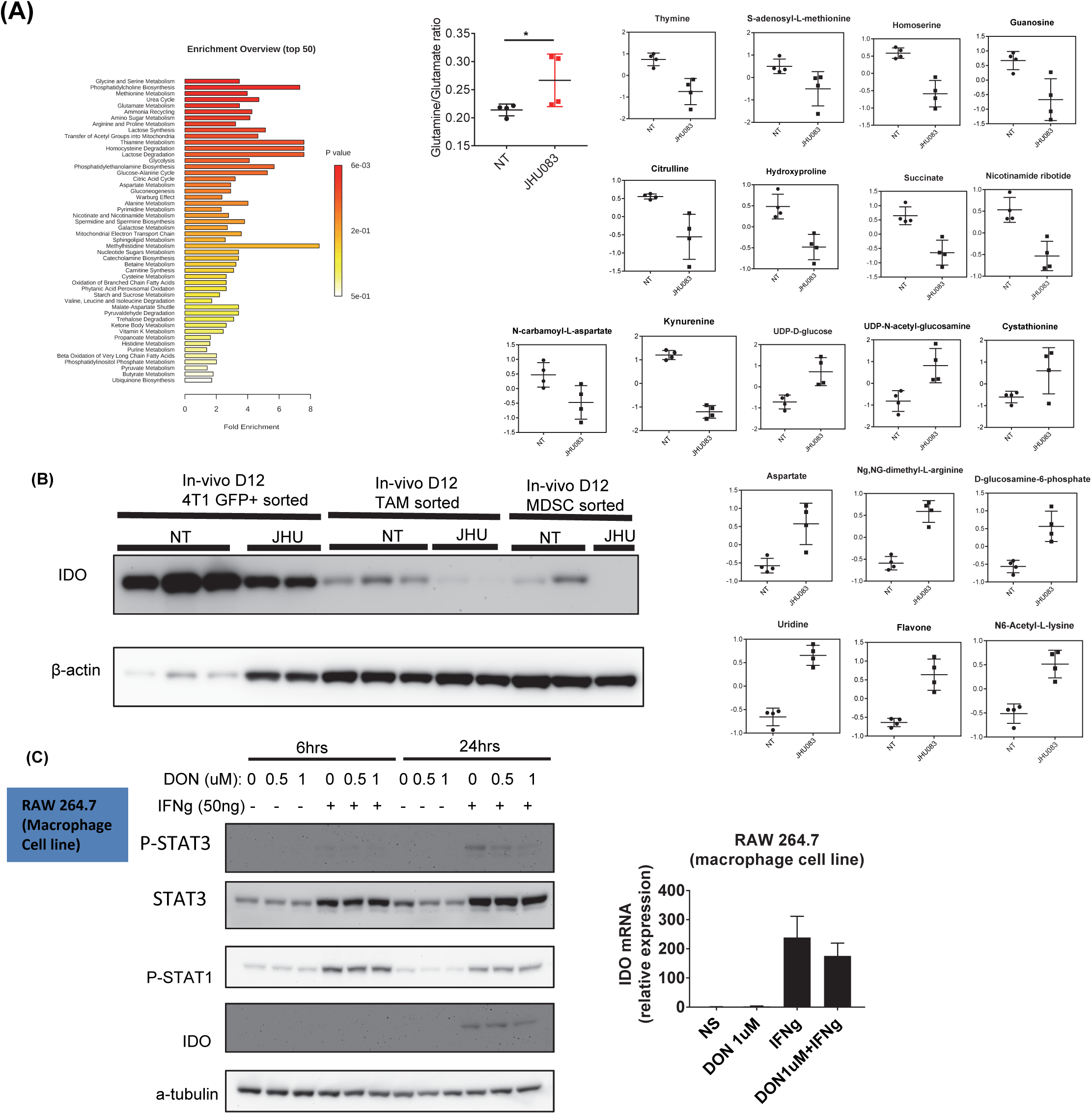
Summary of metabolic changes of tumors from vehicle vs. JHU083 treated 4T1 tumor bearing mice and IDO expression on sorted cells from tumor (A). The pathway association of significantly different metabolites were assessed by pathway enrichment analysis. Graphs of ratio of glutamine and glutamate, and significant metabolites from 4T1 tumors (related to Figure 4A-C). (B) 0.1×10^6^ GFP+4T1 cells were implanted subcutaneously into mammary fat pad of BALB/cJ female mice. 4T1 tumor-bearing mice were treated with JHU083 (1 mg/kg) starting at day 7 after tumor inoculation. After 7 days of treatment, a lower dose (0.3 mg/kg) of JHU083 was used. On day 12, GFP+ tumor cells, TAMs and MDSCs were sorted. Cells were lysed and IDO expression was measured by immunoblot. β-actin was used as loading control. (C) RAW 264.7 cells (macrophage cell line) were cultured in the presence or absence of DON (0.5 μM or 1 μM) and IFNg (50ng) for 6 or 24 hours. P-STAT1 (ser S727), P-STAT3 (Ser727) and IDO1 were measured by immunoblot. α-tubilin were measured as loading controls (left). After 6hrs with DON 1 μM treatment, IDO mRNA expression level was measured by q-PCR (right).

**FigS5 related to Figure 6.**
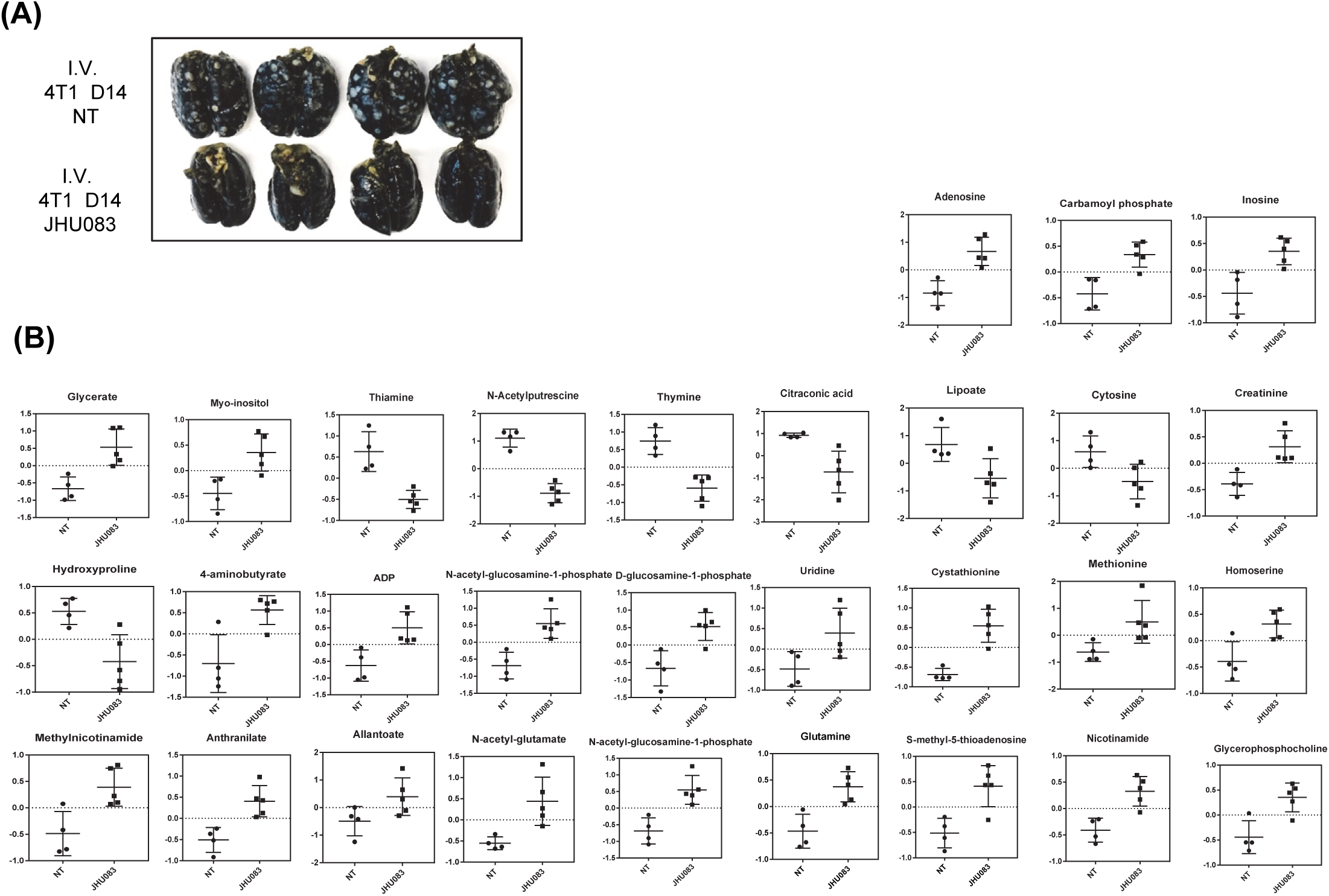
Summary of metabolic changes of lungs from vehicle vs. JHU083 treated 4T1 tumor bearing mice. (A) 0.1×10^6^ 4T1 cells were injected intravenously into the tail vein of Balb/cJ female mice. 4T1 tumor-bearing mice were treated with JHU083 (1mg/kg) starting at day 2 after tumor inoculation. After 7 days of treatment, lower dose (0.3 mg/kg) of JHU083 was used. On day 14, the whole lung were harvested, and lung metastasis were analyzed by inflation with 15% india ink to quantify tumor nodules (B) Significant metabolites from lungs of 4T1 tumor bearing mice (related to Figure 5A-).

**FigS6 related to Figure 7.**
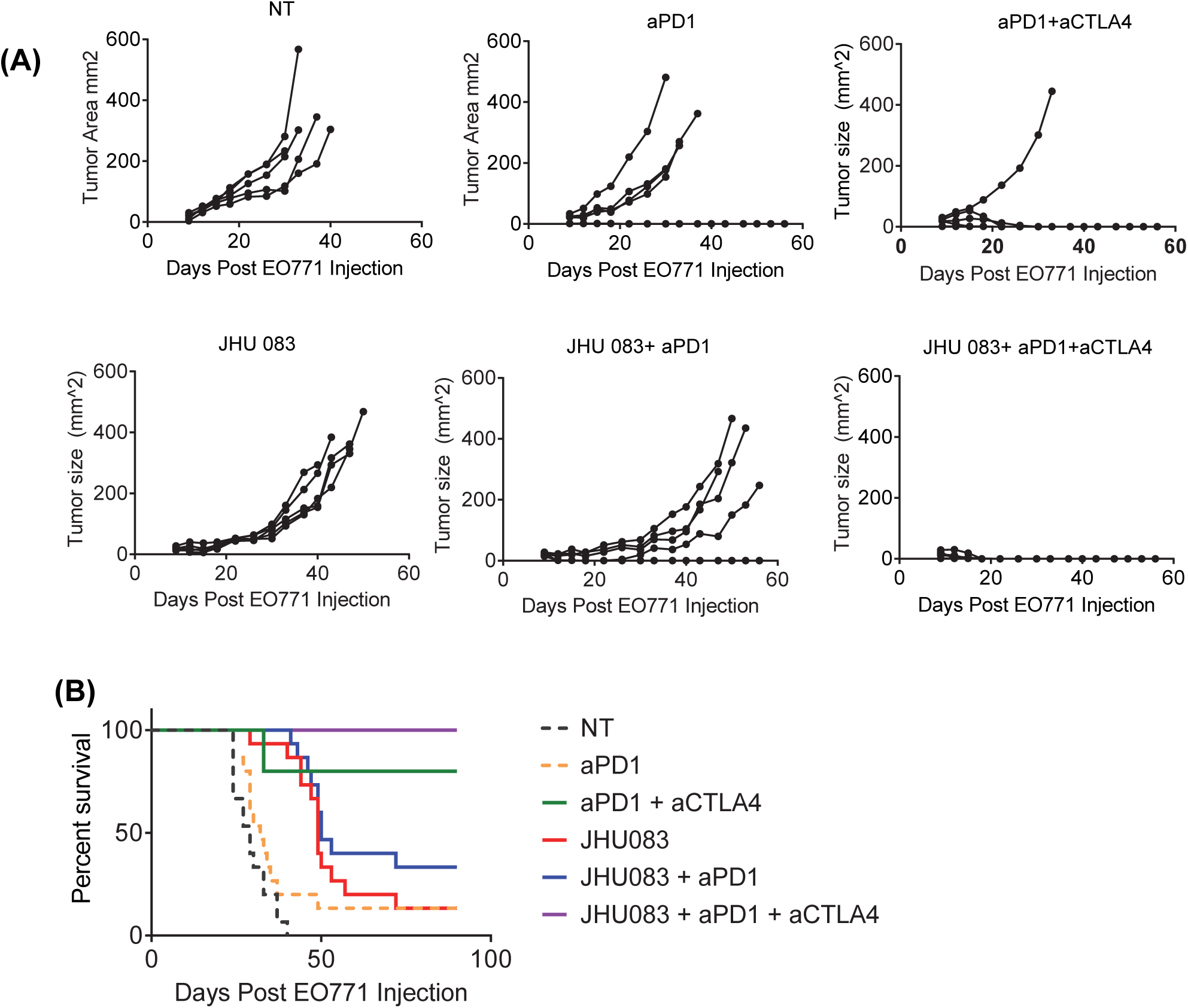
Glutamine antagonism enhances immune checkpoint blockade in EO771 tumor-bearing mice. (A-B) 0.2×10^6^ EO771 cells were implanted subcutaneously into mammary fat pad of C57BL/6J female mice. EO771 tumor-bearing mice were treated with JHU083 (1 mg/kg) daily starting at day 7 after tumor inoculation. After 7 days of treatment, JHU083 daily dose was reduced (0.3 mg/kg). On days 9, 12, and 15, mice were injected with or without 100 μg anti-PD1 followed by treatment with or without JHU083. (A) Each individual mice tumor growth curve and (B) survival curve were recorded (N=5/group). Data are representative of at least three independent experiments. Log-rank (Mantel-Cox) test.

